# psupertime: supervised pseudotime inference for single cell RNA-seq data with sequential labels

**DOI:** 10.1101/622001

**Authors:** Will Macnair, Manfred Claassen

## Abstract

Single cell RNA-seq has been successfully combined with pseudotime inference methods to investigate biological processes which have sequential labels, such as time series studies of development and differentiation. Pseudotime methods developed to date ignore the labels, and where there is substantial variation in the data not associated with the labels (such as cell cycle variation or batch effects), they can fail to find relevant genes. We introduce `psupertime`, a supervised pseudotime approach which outperforms benchmark pseudotime methods by explicitly using the sequential labels as input. `psupertime` uses a simple, regression-based model, which by acknowledging the labels assures that genes relevant to the process, rather than to major drivers of variation, are found. `psupertime` is applicable to the wide range of single cell RNA-seq datasets with sequential labels, derived from either experimental design or user-selected cell cluster sequences, and provides a tool for targeted identification of genes regulated along biological processes.

## 1 Main

Single-cell RNA sequencing studies have been used to define the transcriptional changes corresponding to biological processes, including embryonic development [1], response to stimulus [2], differentiation [3] and aging [4]. Such studies are typically based on single cell RNA-seq measurements of cells representing states of the studied process, often with a sequence of condition labels corresponding to progression along the process, such as timepoints in a time series. Subsequent pseudotime ordering of the individual cells, which computationally orders cells along trajectories, is used to estimate both the state sequence, and genes associated with the process of interest. Many pseudotime techniques have been proposed and are based typically on defining similarities between cells, then identifying an ordering which places similar cells close to each other [5]. These methods assume that the major driver of variation in the data corresponds to the process of interest. However, where there are strong additional sources of variation, these methods are unlikely to identify all relevant genes. For example, batch effects and cell cycle are both known to have strong effects on transcriptional profiles [6, 7]. While techniques to correct for these have been proposed [7–9], even where these techniques are effective, they do not guarantee that the ordering identified by a pseudotime technique will correspond to the condition labels.

To address this problem, we introduce a supervised pseudotime technique, `psupertime`, which explicitly uses sequential condition labels as input (Fig 1A). `psupertime` is based on penalized ordinal logistic regression (Fig 1B), and learns a sparse linear combination of genes which places the cells in the ordering specified by the sequence of labels. This allows for targeted characterization of processes in single cell RNA-seq data, despite substantial variation not associated with the process of interest. Full details of the method are given in **Methods**.

**Figure 1:**
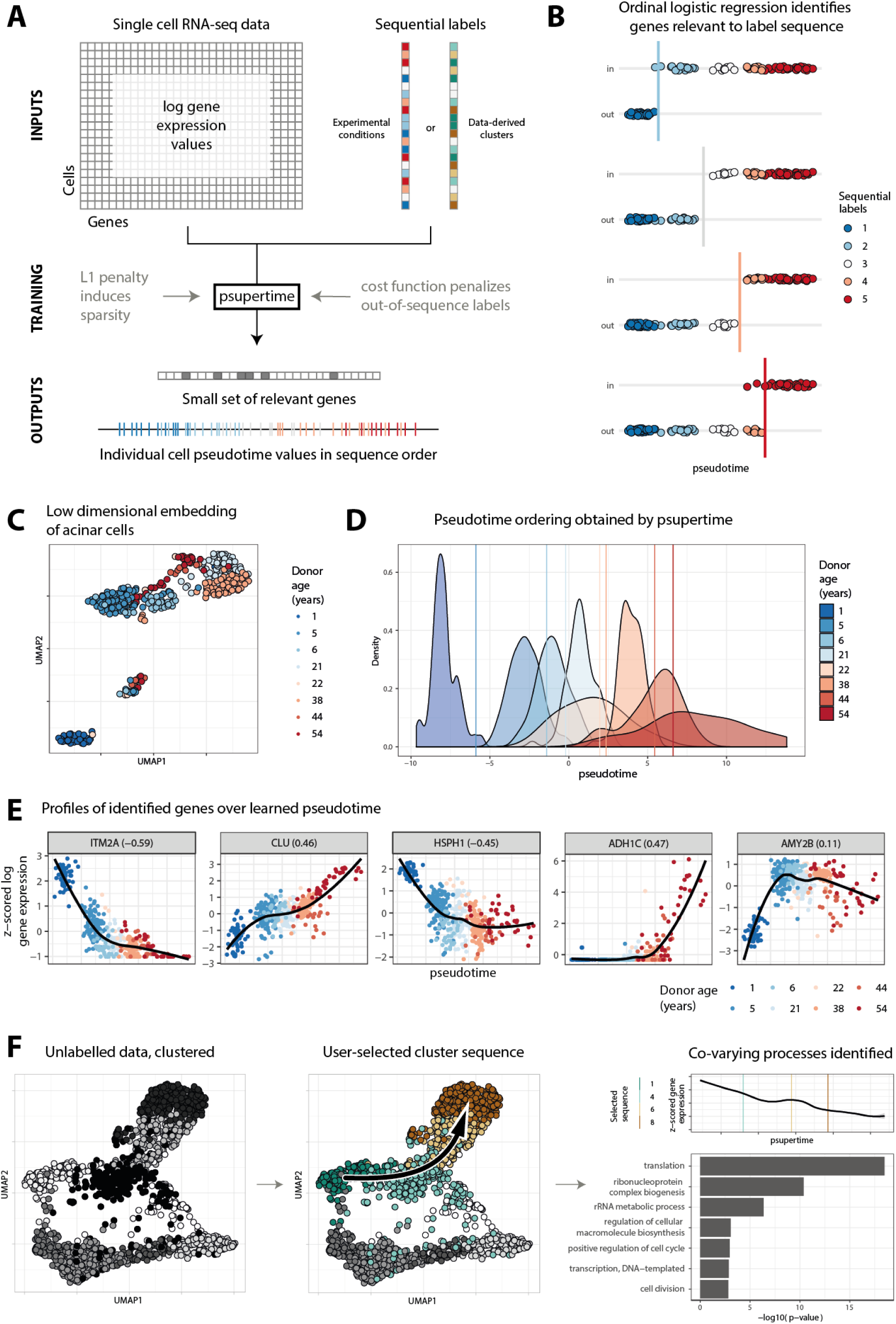
**A** Inputs to `psupertime` are single cell RNA-seq data, where the cells have sequential labels associated with them. `psupertime` then identifies a sparse set of ordering coefficients for the genes. Multiplying the gene expression values by this vector of coefficients gives pseudotime values for each cell, which place the labels approximately in sequence. **B** Cartoon of statistical model used by `psupertime`, including thresholds between labels. Where there is a sequence of *K* condition labels, `psupertime` learns *K* − 1 simultaneous (i.e. sharing coefficients) logistic regressions, each seeking to separate labels 1 *… k −* 1 from *k … K*. **C** Dimensionality reduction of 411 human acinar cell data with ages ranging from 1 to 54 [4]. Representations in two dimensions via non-linear dimensionality reduction technique `UMAP` [21]. Colours indicate donor age. **D** Distributions of donor ages for acinar cells over the pseudotime learned `psupertime`. Vertical lines indicate thresholds learned by `psupertime` distinguishing between earlier and later sets of labels; colour corresponds to the next later label. **E** Expression values of selected genes (five with largest absolute coefficients; see Supp Fig 2 for 20 largest). *x*-axis is `psupertime` value learned for each cell; *y*-axis is z-scored log_2_ gene expression values. Gene labels also show the Kendall’s *τ* correlation between sequential labels (treated as a sequence of integers 1, …, *K*) and gene expression. **F** `psupertime` can be used to explore data without condition labels: unsupervised clustering is first applied to generate labels, then the user selects a sequence of clusters they wish to investigate, and runs `psupertime`, identifying associated genes and processes. Results show dimensionality reduction (`UMAP` [21]) of 1894 colon cells [14], clustered by unsupervised clustering. Plot shows a user-selected sequence of clusters. Hierarchical clustering of gene expression identified 5 gene clusters; illustrative cluster shown here has highest negative correlation with learned pseudotime. Geneset enrichment of clustered gene profiles identifies biological processes associated with this cluster. GO terms shown correspond to the smallest *p*-values, subject to *p <* 0.1% and at least 5 annotated genes. See **Methods** for details, and Supp Results 3 for further analysis.

We demonstrate `psupertime` on a dataset comprising 411 cells from the pancreas, from eight human donors with ages from 1 to 54 years [4]. Acinar cells perform the exocrine function of the pancreas, producing enzymes for the digestive system. This dataset was selected because each set of cells was obtained from different donors, resulting in significant variation in the dataset unrelated to donor age (Fig 1C). Despite this variation, `psupertime` finds a cell-level ordering which respects the age progression, while separating the labels from each other (Fig 1D). We show that the performance of `psupertime` is robust, including to perturbations in labels (see Supp Results 1).

`psupertime` produces as output ordering coefficients for a sparse set of genes, balancing the requirement for predictive accuracy against that for a small and therefore interpretable set of genes. A non-zero ordering coefficient indicates that a gene was was relevant to the label sequence. `psupertime` attains a test accuracy of 83% over the 8 possible labels, using 82 of the 827 highly variable genes (Supp Fig 1). Fig 1E, Supp Fig 2 and Supp Fig 3 show the expression profiles of the genes with highest absolute coefficient values identified along the learned pseudotime. Many of these genes are already known to be relevant to the aging of pancreatic cells: clusterin (*CLU*) plays an essential role in pancreas regeneration, and is expressed in chronic pancreatitis [10, 11]; *α*-amylase (*AMY2B*) is a characteristic gene for mature acinar cells, encoding a digestive enzyme [12]. In addition, `psupertime` suggests candidates for further study: *ITM2A* has the highest absolute gene coefficient, and is highly differentially regulated in a model of chronic pancreatitis, but has not been investigated in acinar cells [13]. The genes identified by `psupertime` were not discussed in the source manuscript, and would not be found by naively calculating correlations between the sequential labels and gene expression (see Supp Results 2).

**Figure 2:**
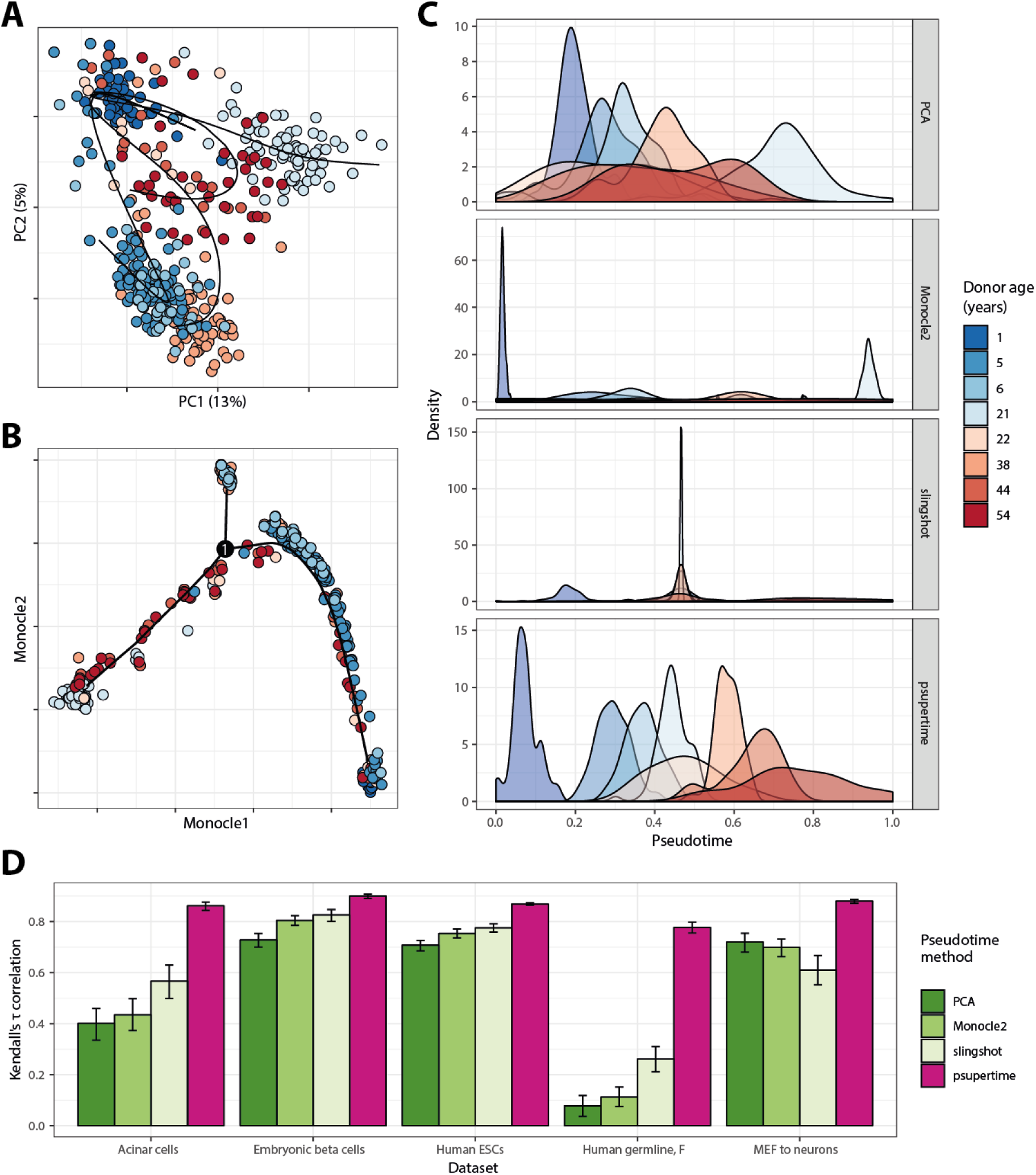
Performance of psupertime against benchmark methods. See subsection 3.5 for details of data processing and use of benchmark methods. All results for **A,B,C** based on 411 aging human acinar cell data with ages ranging from 1 to 54 [4], using 827 highly variable genes. Colours indicate donor age. **A** Projection of acinar cells into first two principal components (% of variance explained shown). Curves learned by slingshot shown (note that here we show the projection of these curves into the first two principal components). **B** Projection of acinar cells into dimensionality reduction calculated by Monocle 2, annotated with pseudotime learned by Monocle 2 [16]. **C** Results of benchmark pseudotime methods applied to acinar data. For each method, the *x*-axis is a one-dimensional representation for each cell (see subsection 3.5), scaled to [0, 1] and given the direction with the highest positive correlation with the label sequence. *y*-axis is density of the distributions for each label used as input, as calculated by the function geom density in the R package ggplot2 [22, 23]. **D** Absolute Kendall’s *τ* correlation coefficient between label sequences (treated as sets of integers 1, … *K*) and calculated pseudotimes. Error bars show 95% confidence interval over 1000 bootstraps, calculated with boot package in R [24]. Datasets specified in Table 1.

**Table 1:** Details of datasets used in benchmark comparisons.

| Dataset name | Source | Accession | Labels used | # labels | # cells | # HVG |
| --- | --- | --- | --- | --- | --- | --- |
| Acinar cells | [4] | GSE81547 | Donor age | 8 | 411 | 827 |
| Human germline, F | [18] | GSE86146 | Age (weeks) | 12 | 992 | 1081 |
| Embryonic beta cells | [39] | GSE87375 | Developmental stage | 7 | 575 | 2666 |
| Human ESCs | [1] | E-MTAB-3929 | Embryonic day | 5 | 1529 | 2876 |
| MEF to neurons | [2] | GSE67310 | Days since induction | 5 | 315 | 1698 |
| Colon cells | [14] | GSE102698 | User-selected clusters | 4, 5 | 1894 | 1515 |
| iPSCs | [19] | GSE106340 | Days during reprogramming | 11, 9 | 8800 | 837 |

GO term enrichment analysis provides further support for the validity of the cell ordering identified by `psupertime`. We clustered the expression profiles of the highly variable genes, and identified GO terms characteristic of each cluster (see **Methods**). This procedure identified genes related to digestion as being up-regulated in early ages (‘proteolysis’ and ‘digestion’ enriched in cluster 1), and terms related to aging later in the process (‘negative regulation of cell proliferation’ and ‘positive regulation of apoptotic process’ enriched in cluster 5) (see Supp Fig 4, Supp Fig 5). This analysis confirms that the cell ordering learned by `psupertime` is plausible.

In studies where the experimental design does not define sequential condition labels, lower-dimensional embeddings of the data may suggest trajectories within the dataset corresponding to sequences of states traversed by a biological process. To study such trajectories, researchers can specify a sequence of clusters, and use `psupertime` to identify the genes and processes regulated along it. We demonstrate this approach on 1894 unlabelled cells from the colon, where goblet (secreting) and colonocyte (absorbing) cells are known to be renewed by stem cells [14] (Fig 1F). The dimensionality reduction embedding shows subpopulations of similar cells, derived via unsupervised clustering [15] (see **Methods**), and suggests possible trajectories of interest that can be defined in terms of these clusters. `psupertime` was applied to the cluster sequence shown. Combined with clustering of genes and GSEA, this analysis indicates that the state sequence corresponds to stem cells which differentiate into colonocytes (genes up-regulated early in the process are related to translation and cell division, while those expressed later in the process correspond to transport; see Supp Results 3). Such datasets are typically analysed by applying unsupervised pseudotime inference techniques, however this does not allow users to specify the sequence of cell clusters they wish to investigate. `psupertime` therefore provides a method for targeted exploratory data analysis of unlabelled data, via evaluation of user-defined cell cluster sequences.

We compare `psupertime` to three alternative, unsupervised pseudotime techniques: projection onto the first PCA component, as a simple, interpretable baseline; Monocle 2 [16], which is widely used, shown to perform well in a benchmark study [5] and permits the selection of a starting point; and slingshot [17], which was also shown to perform well [5] and allows both the start and end point of a trajectory to be selected (it is therefore semi-supervised). Applied to the acinar cells, low-dimensional embeddings of the data (including PCA) indicate that while donor-specific factors account for much of the variation, very little transcriptional variation is related to age (Fig 2A,B; Supp Fig 6). Acinar cell orderings identified by the benchmark methods are not consistent with the known label sequence (Fig 2C, Fig 2D). In contrast, the one-dimensional projection learned by `psupertime` (Fig 2C) successfully orders the cells by donor age (Kendall’s *τ* correlation coefficient 0.86, which quantifies the concordance between two orderings), while providing a sparse interpretable gene signature related to age.

In addition to the acinar cells, we compared `psupertime` to the three alternative methods on four further datasets, as specified in Table 1. Performance of the benchmark methods varies considerably depending on the dataset (Table 2), and in particular depending on the extent of variation unrelated to the labels (Supp Fig 6): both Monocle 2 and PCA show Kendall’s *τ* values of 0.12 or below for the human germline dataset [18] (Supp Fig 7), in comparison to values of at least 0.71 for the human ESCs dataset [1] (Supp Fig 8). In all datasets considered, the cell ordering given by `psupertime` has a higher correlation with the known label sequence than the other pseudotime methods (Fig 2D). We note that due to cellular heterogeneity, the known label sequence is an approximation to the pseudotime, i.e. the variable indicating for each cell the progress through the studied process. However, it is the best available estimate of this variable for evaluating the capability of methods to recapitulate the process under investigation, and therefore to identify its relevant genes (see also Supp Results 5).

**Table 2:** Correlations of pseudotimes with known labels, using highly variable genes as input. See subsection 3.5 for details of calculations.

| Dataset | Method | Spearman's $\rho$ | Kendall's $\tau$ |
| --- | --- | --- | --- |
| Acinar cells | PCA | 0.56 | 0.40 |
| Acinar cells | Monocle2 | 0.57 | 0.43 |
| Acinar cells | slingshot | 0.66 | 0.57 |
| Acinar cells | psupertime | 0.96 | 0.86 |
| Human germline, F | PCA | 0.10 | 0.08 |
| Human germline, F | Monocle2 | 0.17 | 0.11 |
| Human germline, F | slingshot | 0.34 | 0.26 |
| Human germline, F | psupertime | 0.91 | 0.78 |
| Embryonic beta cells | PCA | 0.88 | 0.73 |
| Embryonic beta cells | Monocle2 | 0.93 | 0.80 |
| Embryonic beta cells | slingshot | 0.93 | 0.83 |
| Embryonic beta cells | psupertime | 0.98 | 0.90 |
| Human ESCs | PCA | 0.84 | 0.71 |
| Human ESCs | Monocle2 | 0.87 | 0.75 |
| Human ESCs | slingshot | 0.89 | 0.78 |
| Human ESCs | psupertime | 0.97 | 0.87 |
| MEF to neurons | PCA | 0.87 | 0.72 |
| MEF to neurons | Monocle2 | 0.87 | 0.70 |
| MEF to neurons | slingshot | 0.74 | 0.61 |
| MEF to neurons | psupertime | 0.97 | 0.88 |

`psupertime` achieves classification accuracies of between 43% to 98% over the test datasets, and the time taken to run varies from 4s for a dataset with ≈300 cells, to 32s for one with ≈1500 cells (Table 3). `psupertime` achieves its accuracy using a small set of genes: for example, for an accuracy of 76% on the acinar cells, `psupertime` uses 10% of the input genes (Table 3). `psupertime` is based on a form of penalized linear regression. We show that the ordinal logistic model, rather than a linear model based on regarding the sequential labels as integers, is both the natural and the best-performing model for this problem (see Supp Results 2).

**Table 3:** psupertime performance and timings on comparison datasets. Mean and standard deviation of `psupertime` accuracy, timing and sparsity calculated over 10 random seeds.

| Dataset name | Accuracy (%) | Time taken (s) | Sparsity (%) |
| --- | --- | --- | --- |
| Acinar cells | 75.7 ± 1.1 | 5.5 ± 0.41 | 90.6 ± 1.8 |
| Human germline, F | 43.4 ± 1.5 | 25 ± 0.73 | 80.4 ± 4.9 |
| Embryonic beta cells | 78.5 ± 0.9 | 19 ± 0.67 | 96.4 ± 0.4 |
| Human ESCs | 97.6 ± 0.2 | 35 ± 0.80 | 90.0 ± 1.1 |
| MEF to neurons | 89.6 ± 1.7 | 4.7 ± 0.062 | 96.6 ± 0.4 |

Typical workflows for single cell RNA-seq data first restrict to highly variable genes. If the data is instead first restricted to genes which correlate strongly with the sequential labels, the performance of the benchmark methods might become competitive. Despite the selection of genes that correlate with the labels, `psupertime` consistently outperforms the unsupervised methods (Supp Results 2). This illustrates that the genes identified by `psupertime` as most relevant to the process are not necessarily those with highest correlation; for example, genes with expression profiles like *AMY2B* in Fig 1E show a non-linear, step-like expression profile, which results in a correlation of 0.11 with the condition labels. Despite low correlation, such genes were nonetheless found to be useful for cell ordering, and suggest that `psupertime` discovers meaningful non-linear structure in the data.

Once `psupertime` is trained, it can then be used to predict labels for new data with different or unknown labels. Specifically, we evaluated how the course of induced pluripotent stem cell (iPSC) reprogramming is affected by different culture conditions Schiebinger *et al.* We applied `psupertime` to mouse embryonic fibroblast cells (MEFs) treated to become iPSC, under two different protocols both known to produce iPSCs: treatment with either fetal bovine serum (‘serum’), or with a combination of two inhibitors (‘2i’). Training `psupertime` on cells under the serum condition, and predicting the pseudotime values for cells from the 2i condition, indicates that even the fully mature cells treated with serum are only equivalent to day 11 or 12 with respect to the reprogramming process under 2i (Supp Fig 9, Supp Fig 10). This demonstrates that `psupertime` can be used to assess cells obtained under one process with respect to another.

The number of studies using single cell RNA-seq is increasing exponentially [5]. Many of these include cell groups annotated with condition labels, where the labels have a natural sequence, such as the timepoints of a time series experiment. `psupertime` is explicitly designed to take advantage of such a setting, in contrast to unsupervised pseudotime techniques. The presence of condition labels allows a simple, regression-based model to outperform the more sophisticated pseudotime approaches required for unlabelled data. It efficiently discriminates between relevant and unwanted variation, enabling it to identify genes which recapitulate the ordering, even where batch or other confounding effects prove too challenging for unsupervised techniques. `psupertime` is applicable to any experimental design with sequential labels, most obviously time series but also to biological questions regarding drug dose-response, and disease progression. We have described several additional uses for `psupertime`: exploration of data without experimental labels by combining with unsupervised clustering analysis (see Supp Results 3); alignment of new data to orderings learned from alternative processes (see Supp Results 3). More broadly, we have used it to improve dimensionality reduction (see Supp Results 4), and are developing extensions including to additional single cell technologies such as mass cytometry [20] (see Supp Results 5). This demonstrates the potential of ordinal regression models for further methodological developments. `psupertime` has wide applicability, and will enable quick and effective identification of the genes and profiles relevant to state sequences of biological processes in single-cell RNA sequencing data. We have developed an R package available for download at github.com/wmacnair/psupertime.

## 3 Methods

### 3.1 Overview of psupertime methodology

`psupertime` requires two inputs: (1) a matrix of log read counts from single cell RNA-seq, where rows correspond to genes and columns correspond to cells; and (2) a set of labels for the cells, with a defined sequence for the labels (for example, a set of cells could have labels *day1*, *day3*, *day1*, *day2*, *day3*). (Note that not all cells need to be labelled: `psupertime` can also be run on a labelled subset.) `psupertime` then identifies a set of ordering coefficients, *β_i_*, one for each gene (Fig 1A). Multiplication by this vector of coefficients converts the matrix of log gene expression values into pseudotime values for each individual cell. The set of pseudotime values recapitulates the known label sequence (so the cells with labels *day1* will on average have lower pseudotime values than those labelled *day2*, and so on). The vector of coefficients is *sparse*, in the sense that many of the values are zero; these therefore have no influence on the ordering of the cells. Genes with non-zero coefficients are therefore identified by `psupertime` as relevant to the process which generated the sequential labels.

Suppose the sequence of condition labels we have is 1, …, *K*. Intuitively, `psupertime` learns a weighted average of gene expression values that separates the cells with label 1 from the cells with labels 2, …, *K*, at the same time as separating 1, 2 from 3, …, *K*, and 1, 2, 3 from 4, …, *K*, and so on (Fig 1B). This can be thought of as solving *K* − 1 simultaneous logistic regression problems, and is termed *ordinal logistic regression* [25].

As described so far, `psupertime` can be thought of as minimizing a cost, where the cost is the error in the resulting ordering. To make the results more interpretable, we would like `psupertime` to use a small set of genes for prediction. To do this, we add a cost for each coefficient *β_i_* used, so that `psupertime` is minimizing *error* + *λ*Ʃ_*i*_ |*β_i_*|; approaches like this are termed ‘regularization’, and in this case ‘L1 regularization’. The parameter *λ* controls the balance between minimizing error, and minimizing the ‘coefficient cost’. The method for implementing this approach is based on the R package glmnetcr, which we have extended with an additional statistical model.

The results of this procedure are: (1) a small and therefore interpretable set of genes with non-zero coefficients; (2) a pseudotime value for each individual cell, obtained by multiplying the log gene expression values by the vector of coefficients; and (3) a set of values along the pseudotime axis indicating the thresholds between successive sequential labels (these can then be used for classification of new samples). Where the data does not have condition labels, `psupertime` can be combined with unsupervised clustering to identify relevant processes (see Supp Results 3). `psupertime` also includes a set of functions for plotting the results.

### 3.2 Pre-processing of data

To restrict the analysis to relevant genes and denoise the data, `psupertime` first applies pre-processing to the log transcripts per million (TPM) values. Specifically, `psupertime` first restricts to highly variable genes, as defined in the scran package in R, i.e. genes that show above the expected variance relative to genes with similar mean expression [26]. Genes that are only expressed in a small number of cells (the default is 1%) are excluded. (The default in `psupertime` is to select highly variable genes. However, prior knowledge regarding the underlying biological process can also be reflected, by selecting a known set of genes for input, such as transcription factors or selected GO terms.)

Single cell RNA-seq data is known to be noisy, and in particular to suffer from dropout, i.e. zero read count values that are technical artefacts [27]. A simple approach to address this is denoising of the cells, by replacing the values at a given cell with an average of its neighbours. `psupertime` implements this by calculating correlations between the log expression values across all selected genes for each pair of cells, using the correlations to identify the 10 nearest neighbours for each cell, and replacing the value for a given cell by the mean value over these neighbours. This step also addresses dropouts. There are many alternative procedures for denoising and dropout imputation in single cell RNA-seq data [28–31]; users may choose their preferred processing approach before running `psupertime`.

Finally, the resulting log-count values for each gene are scaled to have mean zero and standard deviation one; this step is important, as it ensures that selection of genes is not biased towards those with higher expression or variance.

### 3.3 Penalized ordinal logistic regression

`psupertime` applies cross-validated regularized ordinal logistic regression to the processed data, using the labels as the sequence. Simple logistic regression regresses onto a binary outcome variable, learning a linear combination of input variables that separates the two labels. Ordinal logistic regression is an extension of this to an outcome variable with more than two labels, where the labels have a known or hypothesized sequence. The likelihood for ordinal logistic regression is defined by multiple simultaneous logistic regressions, where each one models the probability of a given observation having an earlier or later label, with the definition of ‘early’/‘late’ differing across the simultaneous regressions (Fig 1B). The same linear combination of input variables is used across all individual logistic regressions. This specific model of ordinal logistic regression, in which the simultaneous logistic regressions each seek to separate labels 1 … *k* from labels *k* + 1 … *K*, is termed *proportional odds*. (A commonly used alternative is the *continuation ratio* model, where the regressions seek to separate labels 1 … *k* from label *k* + 1 alone. This is also implemented as an option in `psupertime`.)

In the case where the number of input variables is high relative to number of observations and may include many uninformative variables, as is common in single cell RNA-seq, it can be helpful to introduce sparsity (i.e. to increase the number of zero coefficients). `psupertime` uses *L1 regularization* to do this [32]. Our approach is based on that in the R package `glmnetcr` [33], which reformulates the data and associated likelihood functions into one single regression model, to take advantage of the fast performance of the `glmnet` package [34]. The model originally implemented in `glmnetcr` is the continuation ratio likelihood; we have extended this approach to implement the proportional odds likelihood, as this model is more appropriate for assessing an entire biological process.

Given input data X ∈ ℝ^*n*×*p*^ and *y* ∈ ℕ^n^ condition labels (which for simplicity we are assume are integers), this results in the following cumulative distribution function for ordinal logistic regression:

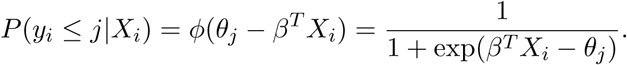

Here, *X_i_* and *y_i_* are the vector and integer corresponding to the *i*th observation and label respectively, *β* is the vector of coefficients and {*θ_j_*} are the thresholds between labels. *ϕ* is the logit link function, which transforms the linear combination of predictors into a probability. Note that the probability given here is cumulative, and that to calculate the probability of an individual label, we have to calculate the difference between successive labels. This results in the following *unpenalized* likelihood:

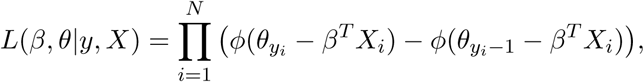

where *y_i_* is the label of observation *i*. Including the L1 penalty, for a given value of *λ*, we obtain the optimal values of *β* and *θ* by maximizing the following penalized objective function:

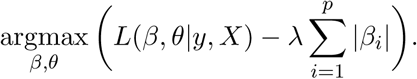

`psupertime` uses cross-validation (with 5 folds as default) to identify the optimal level of L1 regularization: the optimal *λ* is the value with the highest mean score over all held-out folds (either accuracy or cross-entropy may be selected as the score; the default is cross-entropy). To increase sparsity, we use the highest value of *λ* with mean training score within one standard error of the optimal *λ*, rather than take the optimal *λ* itself (following [34]). The model is then retrained using all training data, with this value of *λ*, to obtain the best fitting model.

Where `psupertime` is used to classify completely new data (e.g. from a different experiment), to make the predictions more robust, the cross-validation should take data structure into account (for example, selecting entire samples to be left out, rather than cells selected at random).

### 3.4 psupertime outputs

The `psupertime` procedure results in a set of coefficients for all input genes (many of which will be zero) that can be used to project each cell onto a pseudotime axis, and a set of cutoffs indicating the thresholds between successive sequential labels (Fig 1D). These can be analysed in various useful ways.

The small, interpretable set of genes reported to have non-zero coefficients permits both validation that the procedure has been successful (by observation of genes known to be relevant to the process) and discovery of new relevant genes. The magnitude of a coefficient is a measure of the contribution of this gene to the cell ordering. More precisely, for a gene *i* with coefficient *β_i_*, each unit increase in log transcript abundance multiplies the odds ratio between earlier and later labels by *e^βi^*. Where *β_i_* is small, a Taylor expansion indicates this is approximately equal to a linear increase by a factor of *β_i_*.

The thresholds indicate the points along the `psupertime` axis at which the probability of label membership is equal for labels before the cutoff, and after the cutoff. The distances between thresholds, namely the size of transcriptional difference between successive labels, is not assumed to be constant, and is learned by `psupertime`. Distances between thresholds therefore indicate dissimilarity between adjacent labels, and thresholds which are close together suggest labels which are transcriptionally difficult to distinguish.

The learned geneset can also be used as input to dimensionality reduction algorithms such as t-SNE [35] or UMAP [21]; this is discussed in more detail in Supp Results 4.

Rather than learning a pseudotime for one fixed set of input points, `psupertime` learns a function from transcript abundances to the pseudotime. It can therefore be trained on one set of labels, and applied to new data with unknown or different labels: any data with overlapping gene measurements can be assessed with regard to the learned process. Furthermore, `psupertime` can be learned on two different datasets, with different labels, and then each applied to the other dataset: the sequential labels from one dataset allow coefficients relevant to that sequence to be learned, which can then be used to predict these labels for the second dataset. Seefor more discussion.

### 3.5 Comparison with benchmark methods

To compare the methods, we first performed common preprocessing and identification of relevant genes for each dataset, to identify either highly variable genes, or genes showing high correlation with the label sequence. See Supp Results 2 for further discussion.

To identify highly variable genes, we followed the procedure described by Lun *et al.*, using an FDR cutoff of 10% and biological variability cutoff of 0.5 (see [26] for details of these parameters). To identify genes showing high correlation with the labels, we calculated the Spearman’s correlation coefficient between sequential labels converted into integers, and log gene expression value. Genes with absolute correlation > 0.2 were selected.

For PCA, we calculated the first principal component of the log counts, and used this as the pseudotime. Calculation of `Monocle2` uses the following default settings: genes with mean expression < 0.1 or expressed in < 10 cells filtered out; `negbinomial` expression family used; dimensionality reduction method *DDRTree*; root state selected as the state with highest number of cells from the first label; function `orderCells` used to extract the ordering. Calculation of `slingshot` uses the following default settings: first 10 PCA components used as dimensionality reduction; clustering via Gaussian mixture model clustering using the R package `mclust` [36], number of clusters selected by Bayesian information criterion; root and leaf clusters selected as the clusters with highest number of cells from the earliest and latest labels respectively; lineage selected for pseudotime is path from root to leaf cluster. **Note:** For cells very distant from the selected path, `slingshot` does not give a pseudotime value. For these cells, we assigned the mean pseudotime value over those which slingshot did calculate. Calculation of `psupertime` used default settings, as described in section 3.

We tested the extent to which each pseudotime method could correctly order the cells by calculating measures of correlation between the learned pseudotime, and the sequential labels. Kendall’s *τ* considers pairs of points, and calculates the proportion of pairs in which the rank ordering within the pair is the same across both possible rankings.

To identify genes with high correlation with the sequential condition labels (Table 4), we treated the sequential labels as the set of integers 1, …, *K*, calculated the Spearman correlation coefficient with the gene expression. Genes were selected that showed absolute correlation of > 0.2 with the sequential labels (few genes showed high correlation with the sequential labels; this low cutoff was used to ensure that a sufficient number of genes was selected).

**Table 4:** Correlations of pseudotimes with known labels, using genes correlated with labels as input. See subsection 3.5 for details of calculations.

| Dataset | Method | Spearman's $\rho$ | Kendall's $\tau$ |
| --- | --- | --- | --- |
| Acinar cells | PCA | 0.68 | 0.54 |
| Acinar cells | Monocle2 | 0.73 | 0.58 |
| Acinar cells | slingshot | 0.61 | 0.53 |
| Acinar cells | psupertime | 0.94 | 0.82 |
| Human germline, F | PCA | 0.51 | 0.39 |
| Human germline, F | Monocle2 | 0.49 | 0.36 |
| Human germline, F | slingshot | 0.49 | 0.40 |
| Human germline, F | psupertime | 0.87 | 0.72 |
| Embryonic beta cells | PCA | 0.90 | 0.76 |
| Embryonic beta cells | Monocle2 | 0.92 | 0.79 |
| Embryonic beta cells | slingshot | 0.93 | 0.82 |
| Embryonic beta cells | psupertime | 0.98 | 0.90 |
| Human ESCs | PCA | 0.81 | 0.67 |
| Human ESCs | Monocle2 | 0.89 | 0.77 |
| Human ESCs | slingshot | 0.48 | 0.40 |
| Human ESCs | psupertime | 0.97 | 0.87 |
| MEF to neurons | PCA | 0.86 | 0.71 |
| MEF to neurons | Monocle2 | 0.89 | 0.73 |
| MEF to neurons | slingshot | 0.74 | 0.62 |
| MEF to neurons | psupertime | 0.97 | 0.88 |

### 3.6 Identification of relevant biological processes

To identify biological processes associated with the condition labels, `psupertime` first clusters all genes selected for training (e.g. the default highly variable genes), using the R package `fastcluster` [37], using 5 clusters by default. These are ordered by correlation of the mean expression values with the learned pseudotime, i.e. approximately into genes which are up- or down-regulated along the course of the labelled process. `psupertime` then uses `topGO` to identify biological processes enriched in each cluster, relative to the remaining clusters; enriched GO terms are calculated using algorithm=‘weight’ and statistic=‘fisher’ [38].

## Author contributions

WM conceived the method, wrote the code, devised and performed the analysis, and wrote the manuscript. MC wrote the manuscript.

## 6 Supplementary Results

### Supplementary Results 1: psupertime is robust to label perturbations

The set of condition labels used to train `psupertime` is critical to both the geneset learned by `psupertime`, and the accuracy which can be achieved. If the condition labels progress at a different rate to the underlying biological process, the flexibility in defining thresholds between labels means that `psupertime`’s performance should not be affected. However, if some of the labels are mislabelled, this may be reflected in reduced classification accuracy for a subset of the labels.

We demonstrate that the performance of `psupertime` is robust to a number of perturbations which are plausibly relevant to analysis. We measured the test accuracy of `psupertime` performance under the following perturbations:

1. using different random seeds for the cross-validation folds;
2. selecting at random a pair of neighbouring labels, and swapping their order;
3. completely randomizing the label order; and
4. randomizing the labels of all cells.

We analysed `psupertime`’s performance on the datasets detailed in Table 1. For each dataset, we first restricted to highly variable genes, using the default settings for `psupertime`. For each of the perturbations, we ran `psupertime` twenty times and recorded relevant performance measures: test classification error, test cross-entropy, and sparsity.

The performance of `psupertime` is robust to choice of cross-validation fold, resulting in only small variations in test classification error (Supp Fig 11). (We note that classification error is a volatile measurement, as it is based on 10% of a relatively small number of cells.) Swapping pairs of neighbouring labels results in a small increase in `psupertime` classification error, indicating that small experimental flaws or perturbations do not result in substantial reduction in performance for other labels.

Where the order of labels is randomized, the performance of `psupertime` is reduced, but for some datasets this reduction was small. This may indicate that within the large number of highly variable genes used as input (between ≈ 800 and ≈ 2900; see Table 1), there are sufficient genes to recapitulate a given order of a relatively small number of labels, for a relatively small number of observations.

The number of non-zero genes required to achieve a given level of classification error (i.e. the sparsity) is more variable. Where the cell labels are completely randomized, `psupertime` consistently identifies no genes as being relevant to the ordering, showing that it does not find spurious genes where there is no structure to the data.

The perturbations discussed here correspond to potential mislabelling of the data, which could pose a challenge to `psupertime`. A further challenge comes from data containing branching structure, in which progress along the biological process is accompanied by a bimodal (or multimodal) distribution of expression for some genes. This results in increased variance in gene expression, which could make pseudotime inference more difficult. The cells measured in Schiebinger *et al.* undergo differentiation into multiple distinct celltypes, including iPSCs (see Figure 1 in [19]). Applied to these cells, `psupertime` achieved accurate recapitulation of the label sequence, despite bimodal gene expression for later labels resulting from branching (e.g. *Dppa5a*, *Tmem176b* in Supp Fig 12).

`psupertime` could be affected by the presence of cell sub-populations unrelated to the sequential labels. If this population is consistent across the labels, we expect that `psupertime` would identify relevant genes, although it would have lower accuracy due to being unable to accurately place the unrelated cells. Where there are variable unrelated subpopulations, filtering them out before applying `psupertime` should improve performance. Variability in cell population which is *related* to the sequential labels is not expected to affect the performance of `psupertime`.

Taken together, these results indicate that the performance of `psupertime` is robust, in particular to perturbations in the labels used.

### Supplementary Results 2: psupertime outperforms benchmark methods irrespective of gene selection method

Single cell RNA-seq datasets include measurements of thousands of genes, or features, many of whose measurements may be noisy, or irrelevant to the processes generating the dataset. To identify relevant features, a subset of genes is selected according to statistical criteria. The default criteria implemented in `psupertime` are those proposed by Lun *et al.* and implemented in the R package scran: they note a consistent relationship between the mean and variance of log gene expression data, and select genes which show above average variance given their mean expression. Such an approach is useful to identify relevant genes in an unbiased way, however it does not take into account the information given by sequential labels. Selecting genes which co-vary with the sequential labels should restrict the data to a subset relevant to the labelled process, and not just the genes varying over the dataset (which might therefore include genes affected by batch effects, for example). Co-varying genes can be selected by calculating correlation values, or more generally by identifying genes where a significant proportion of expression variance is explained by the labels (e.g. via ANOVA).

By restricting to genes co-varying with the process, selecting such genes could in principle affect the results of comparisons between `psupertime` and unsupervised methods. We considered four approaches for selecting relevant genes:

1. highly variable genes, following Lun *et al.* [26] and using selection criteria FDR < 0.10, biological component > 0.5;
2. treating the sequential labels as integers 1, …, *K*, and selecting genes with absolute Spearman correlation > 0.2;
3. treating the sequential labels as integers 1, …, *K*, and selecting genes with absolute Kendall’s *τ >* 0.2; and
4. performing ANOVA on all genes, using the labels as the group variable, and selecting genes with *p <* 1e-20, and standard deviation > 2.

For each dataset, we identified genes on the basis of these criteria, and used these as input into `psupertime` and the comparator methods. The Kendall’s *τ* correlation was calculated between the identified pseudotimes and the sequential labels, treated as integers (see subsection 3.5 for details).

The relative performances of `psupertime` and the benchmark methods remain broadly the same across the different methods of gene selection. In particular, under all methods of gene selection, and across all datasets, `psupertime` attains higher correlations than the unsupervised methods (Supp Fig 13). This indicates that even after selecting genes which co-vary with the sequential labels denoting the process of interest, it is necessary to use these labels directly in the inference procedure to obtain pseudotimes which recapitulate the label ordering.

The need for supervised methods is reinforced by the results for LASSO regression [32]. To perform LASSO regression, we converted the sequential labels into integer values 1, …, *K*, and did penalized linear regression. LASSO regression and `psupertime` show similar performance in terms of ability to recapitulate the ordering of the sequential labels, as measured by Kendall’s *τ* (Supp Fig 13), however `psupertime` is better able to classify the cells than LASSO (Supp Fig 14). These results could be expected: treating the sequential labels as integers to be regressed against, as in LASSO, is optimizing for correlation rather than separation, while the thresholds between labels give `psupertime` additional flexibility as a classifier. Taken together, these results suggest that `psupertime` is the appropriate statistical model in terms of both ordering the labels according to the sequence, and accurately labelling the cells.

Comparisons of `psupertime` with unsupervised methods are not strictly fair, as the two types of approach are designed for different tasks: `psupertime` is trained to identify genes covarying with the labels, while the other methods are not. However, unsupervised methods were previously the only methods available for investigating datasets with sequential labels, and have been used for this task; our comparisons are therefore relevant. It is precisely this use of the labels which is the major contribution of `psupertime`.

### Supplementary Results 3: psupertime as a tool for exploratory data analysis of unlabelled single cell RNA-seq data

In studies without sequential condition labels, dimensionality reduction may suggest trajectories within the dataset that are of biological interest. To explore such datasets, researchers can specify a sequence of subpopulations, and use `psupertime` to identify the genes which are regulated along it.

We demonstrate this application on single cell RNA-seq data from the colon, where goblet and colonocyte cells are known to be renewed by stem cells. Supp Fig 15A shows a two-dimensional embedding of 1894 unlabelled cells from the colon [14], indicating several possible trajectories of interest. Unsupervised clustering (Supp Fig 15B) allows trajectories to be specified by the user, two of which are shown in Supp Fig 15C. We used `psupertime`, combined with clustering of genes and geneset enrichment analysis, to identify biological processes characteristic of these trajectories (Supp Fig 15D, Supp Fig 16; see subsection 3.6). Comparison of these results with the discussion in the source manuscript for the data ([14]) suggests that the upper trajectory corresponds to differentiation from stem cells into colonocytes, cells responsible for absorption in the intestine, and that the lower trajectory corresponds to differentiation into goblet cells, which secret mucous (in particular, they express *Muc2*, which had the largest ordering coefficient identified by `psupertime`).

An alternative approach to analysing unlabelled data is to apply unsupervised pseudotime methods, and evaluate the trajectories and co-varying genes they identify. This approach may capture some of the trajectories of interest to users, but users cannot specify exactly which sequence they wish to explore. `psupertime` therefore provides a method for fast, targeted exploration of unlabelled data.

### Supplementary Results 4: psupertime as a feature extraction method for visualization and further analysis

The set of genes reported by `psupertime` correspond to a subset of genes which is relevant to the ordered sequence of labels. This set of genes can therefore be used for feature selection, for example as input to dimensionality reduction algorithms, resulting in a low-dimensional embedding based only on genes specific to the biological process in question.

Restricting to the genes identified as relevant by `psupertime` results in an improvement in the results of dimensionality reduction algorithms (PCA and UMAP [21]), with respect to continuous ordering of the sequential labels (Supp Fig 19). With the exception of the human germline data, all embeddings based on the `psupertime` genes consist of a smaller number of distinct cell clusters than for the HVG genes. The recapitulation of the sequential ordering is also often improved. This is clearest in the cases where the methods already at least partly recapitulated the ordering even with the highly variable genes. Here, restriction to the genes identified by `psupertime` improves the ordering further; for example, applied to the MEF to neurons dataset [2], selecting this subset of genes results in the 20 day old cells being placed in the correct ordering in the PCA plot. Overall, the genes identified by `psupertime` result in embeddings which better reflect the sequential labels, and fewer discontinuities between cells with similar labels.

In principle this feature selection would allow for further analysis of such processes, such as clustering of the cells, or identifying sets of genes showing similar expression profiles. We expect `psupertime` to improve the performance of such methods by excluding genes which are not relevant to the labels.

### Supplementary Results 5: Potential applications and developments of psupertime

We have shown that where sequential labels are available, `psupertime` is able to to identify genes whose expression profiles correspond to this order. It does this even in the presence of substantial unrelated variation, and does so better than benchmark unsupervised methods. `psupertime` is conceptually simple, and its simplicity allows for several avenues for future development.

We performed comparisons between `psupertime` and other unsupervised methods on the basis of ability to recapitulate the ordering of the known labels. Cellular responses are heterogeneous, however, meaning that this known label sequences are imperfect labels of a cell’s progress along a given process. Unsupervised pseudotime techniques explore this heterogeneity by basing their orderings solely on similarities between cells. This suggests a complementary approach between `psupertime` and unsupervised methods: `psupertime` can be used to identify the genes and ordering correlating with the label sequence, and this can be compared with results from unsupervised approaches to quantify the extent of heterogeneity.

While the pseudotime identified by `psupertime` may have a non-linear relationship with the condition labels (for example, in the case where a gene has zero expression for early labels, and a constant higher level of expression for later labels), it is a linear function of the gene expression values. Non-linear implementations of `psupertime` (e.g. as a neural network, or via non-linear regression such as MARS [40]) would allow for pseudotimes which were non-linear, non-monotonic functions of the genes, which would in particular permit the identification of genes showing transient expression.

The design of many single cell studies results in sequential groups of cells (see for example reviews on development [41] and aging [42]). `psupertime` has been developed for single cell RNA-seq data, however it could in principle also be applied to other single cell data such as mass cytometry [20]. We have shown good performance for `psupertime` for single cell RNA-seq data, even though we would not *a priori* expect that a biological process would correspond to a linear combination of gene expressions. This may be the result of the high dimensionality of the dataset, which provides a large set of features with which to approximate a non-linear process. Data derived from mass cytometry is lower-dimensional, and therefore has lower flexibility of marker choice. Here, a non-linear model may be necessary to obtain good performance.

As a classifier, `psupertime` can be first trained on cells with one set of condition labels, then used to project new cells onto the associated process. We showed this by applying `psupertime` to a time course of iPSCs allowed to differentiate, and used this to classify iPSCs kept in pluripotency-maintaining serum (Supp Fig 9). This illustrates potential uses for `psupertime` to compare processes. For example, `psupertime` could be trained on a time series of stimulated cells and used to test the effects of inhibitors: the locations of the inhibited cells, projected onto the pseudotime corresponding to the *uninhibited* process, would indicate the timepoint at which the inhibitor acted.

`psupertime` uses L1 regularization to obtain a small set of reported genes. However, this may result in exclusion of other relevant genes: where there are multiple highly correlated genes that are predictive of the sequential labels, L1 regularization will tend to result in only one of these genes being reported, and produce give zero coefficients for other correlated genes [43]. This issue can be addressed by calculating the `psupertime` ordering, and reviewing all genes that have high correlations with the genes identified by `psupertime`. Alternatively, a trivial extension to `psupertime` would allow training with a combination of L1 and L2 penalties (the *elastic net*), resulting in a compromise between sparsity and prediction performance.

`psupertime` is applicable to many of the increasing number of single cell RNA-seq studies being generated. It shows consistently better performance than benchmark methods, due to use of sequential labels as input. The conceptual novelty of identifying genes via ordinal logistic regression both permits genes relevant to processes annotated with sequential labels to be identified, and suggests new ways of using these labels to understand the genes involved in such processes.

---

**Supp Fig 1:**
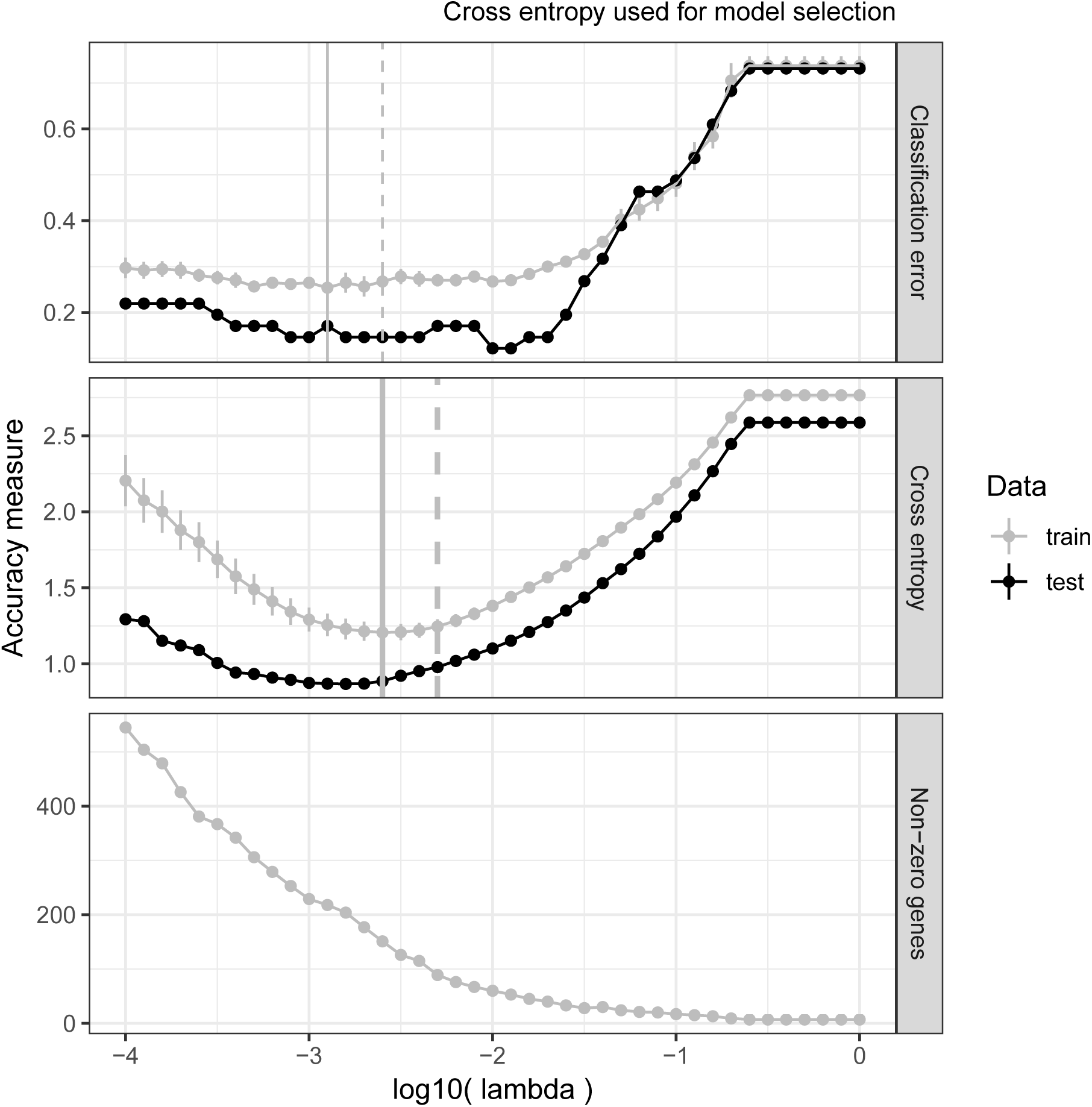
Training and test performance of psupertime applied to acinar cells. *x*-axis is log value of *λ*, the weight given to the *L*_1_ penalty. *y*-axis is a measure of performance. Classification error is the proportion of cells for which `psupertime` predicted the incorrect label. Cross entropy is a measure of how confidently `psupertime` predicted the correct label, and has low values when a correct label is predicted with high probability. Non-zero genes is the number of genes with non-zero coefficients in this model. The grey trend line shows the mean performance measure over the 5 folds in the training data, with vertical whiskers showing the s.e. of the mean. The black trend line shows performance on 10% of data not used for training. Vertical grey lines show the value of lambda with the best performance on the training data (solid) and within one s.e. of the best performance (dashed line); here, the lambda corresponding to the dashed line was selected. The measure used for selection of *λ* (cross-entropy) is indicated with thicker vertical grey lines.

**Supp Fig 2:**
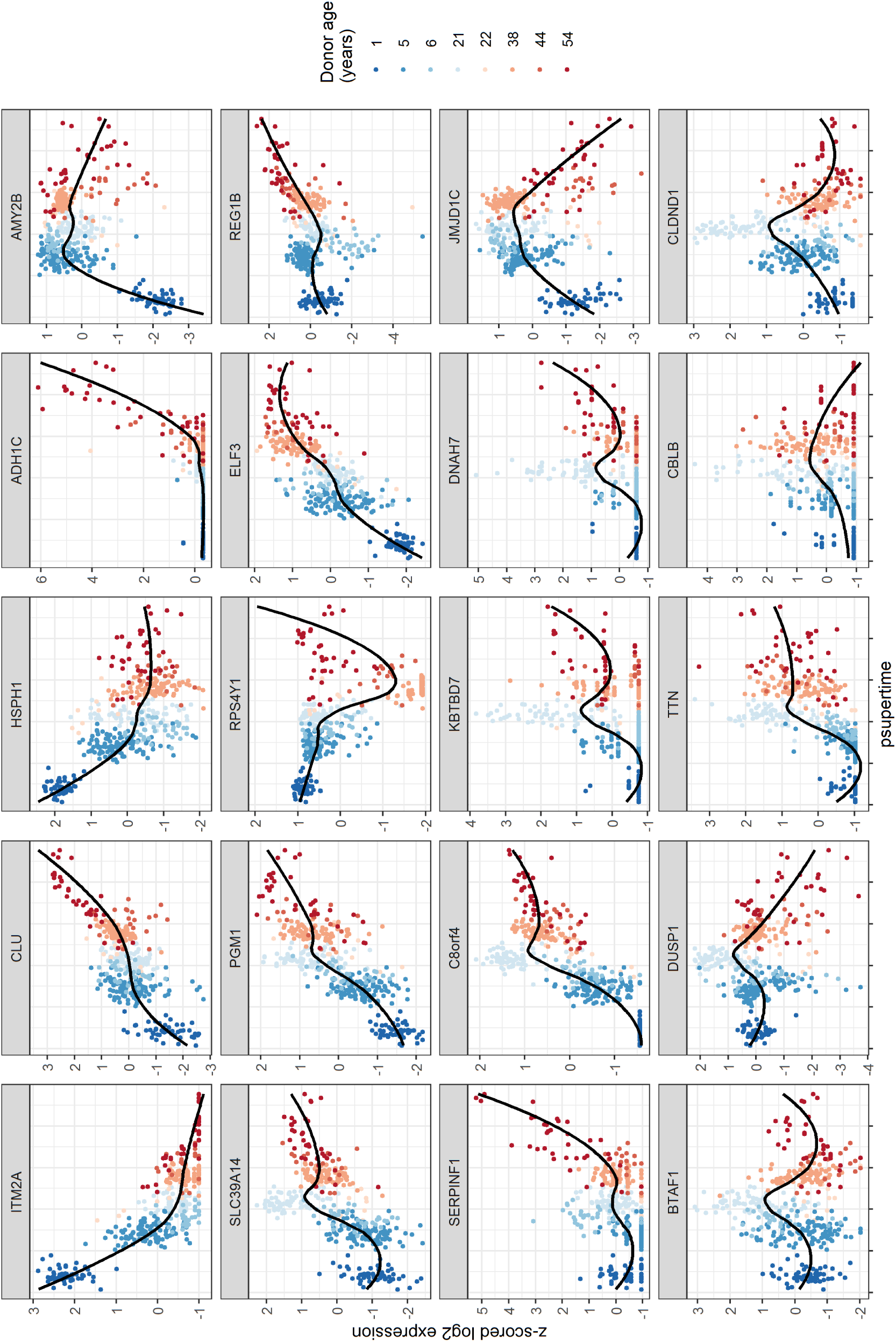
Profiles of top genes identified in acinar cells by psupertime. Results of `psupertime` applied to 411 acinar cells. 20 genes with highest absolute coefficients, plotted against `psupertime` pseudotime. *x*-axis is the values from projections of each cell by `psupertime`. *y*-axis is smoothed, z-scored log pseudocounts for each cell. Colours indicate ordered labels. Black line is smoothed curve as fit by `geom_smooth` in the R package `ggplot2` [23].

**Supp Fig 3:**
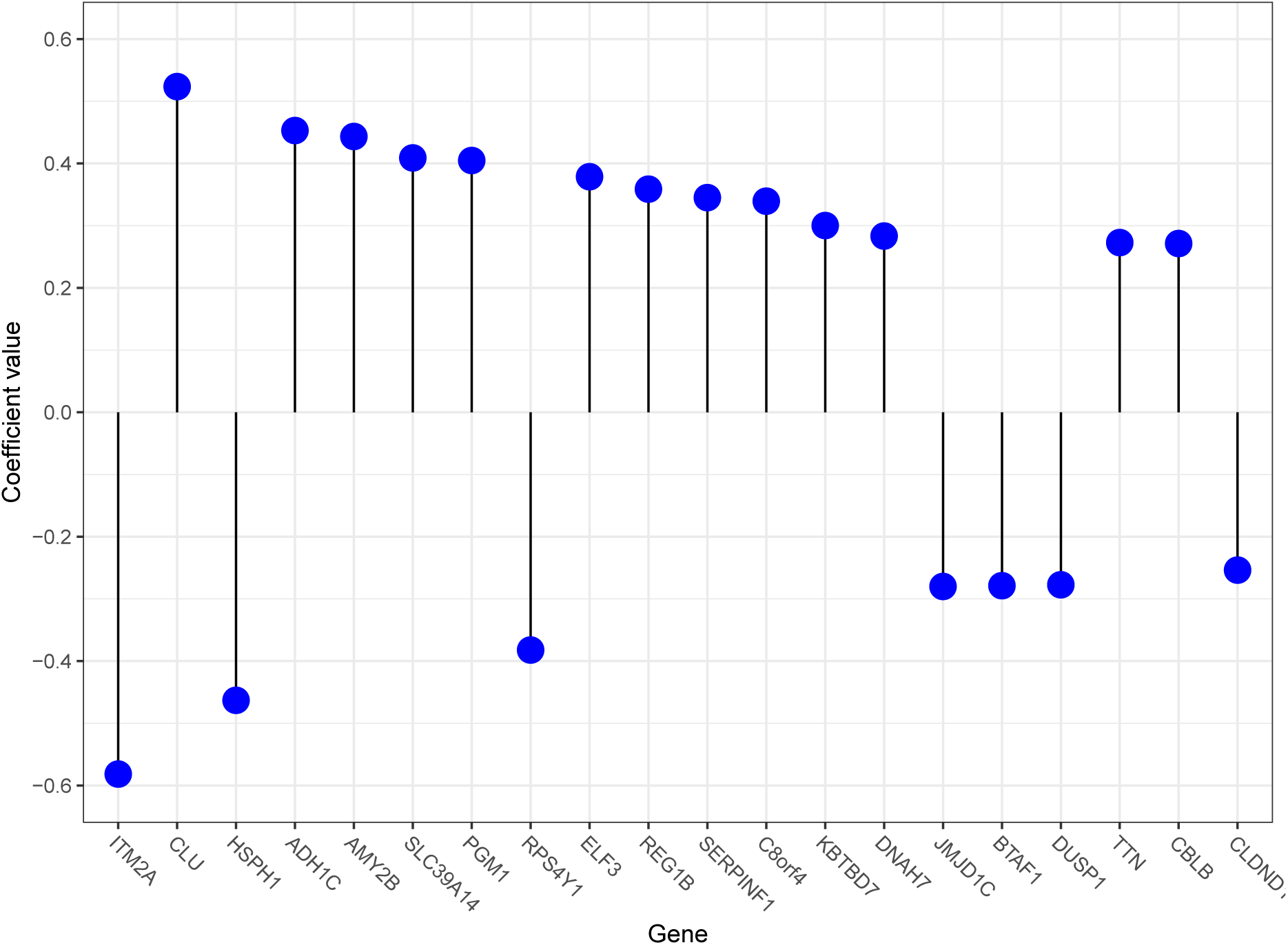
Top genes identified in acinar cells by psupertime. 20 genes with largest absolute ordering coefficients *β_i_*, subject to *β_i_ >* 0.05, ordered by absolute value. Non-zero coefficients correspond to genes relevant to the process underlying the condition labels, and coefficient indicates strength and direction of the effect of this gene on the predicted label.

**Supp Fig 4:**
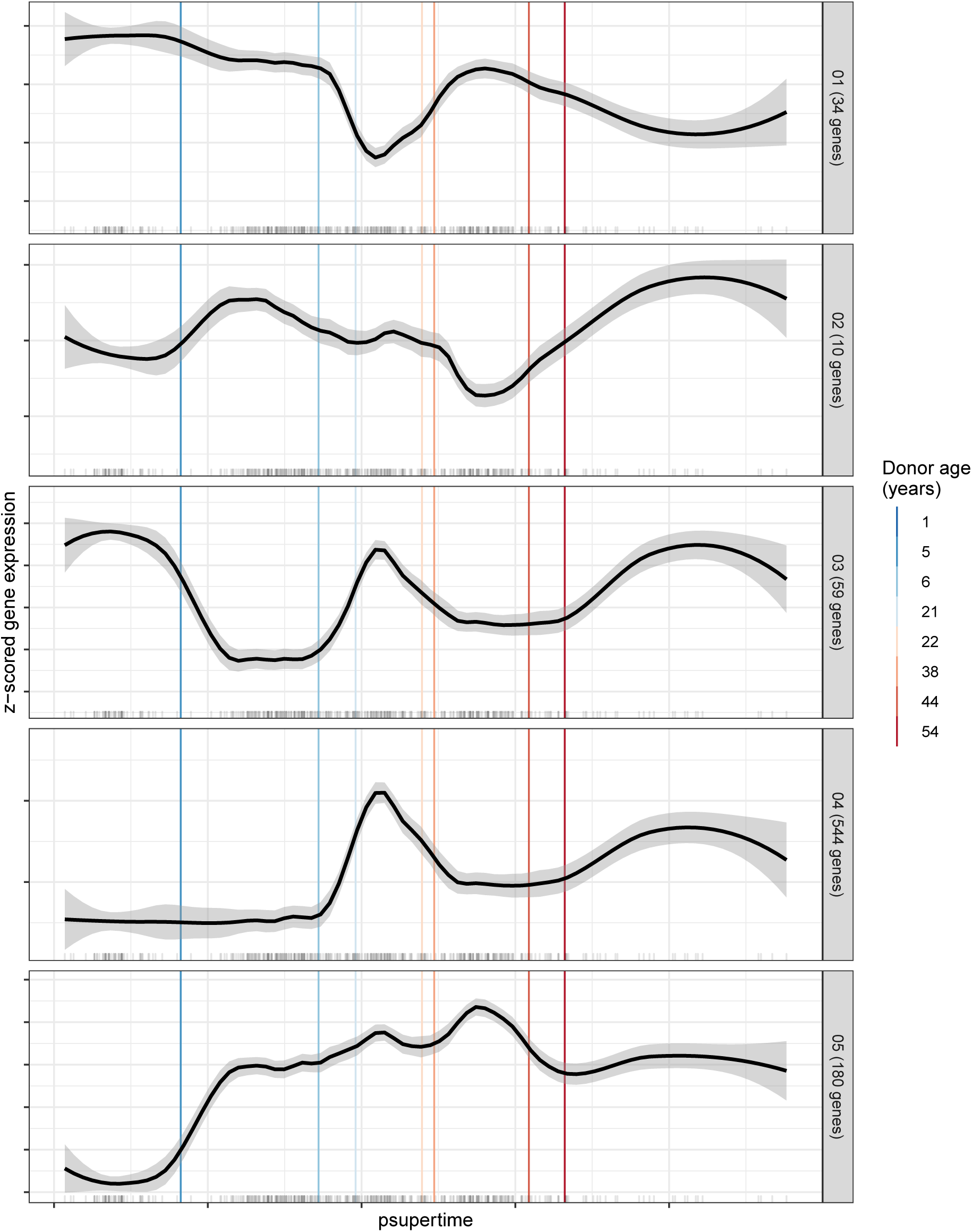
Mean expression profiles of gene clusters, ordered by psupertime. Complete linkage hierarchical clustering was applied to 827 highly variable genes, with *k* = 5 [37]. Black line is smoothed curve as fit by `geom_smooth` in the R package `ggplot2` [23]. Clusters ordered by correlation between mean expression over the gene cluster, and the pseudotime values learned by `psupertime`. Vertical lines indicating predicted cutoffs separating labels. Marks on *x*-axis denote pseudotime values of individual cells.

**Supp Fig 5:**
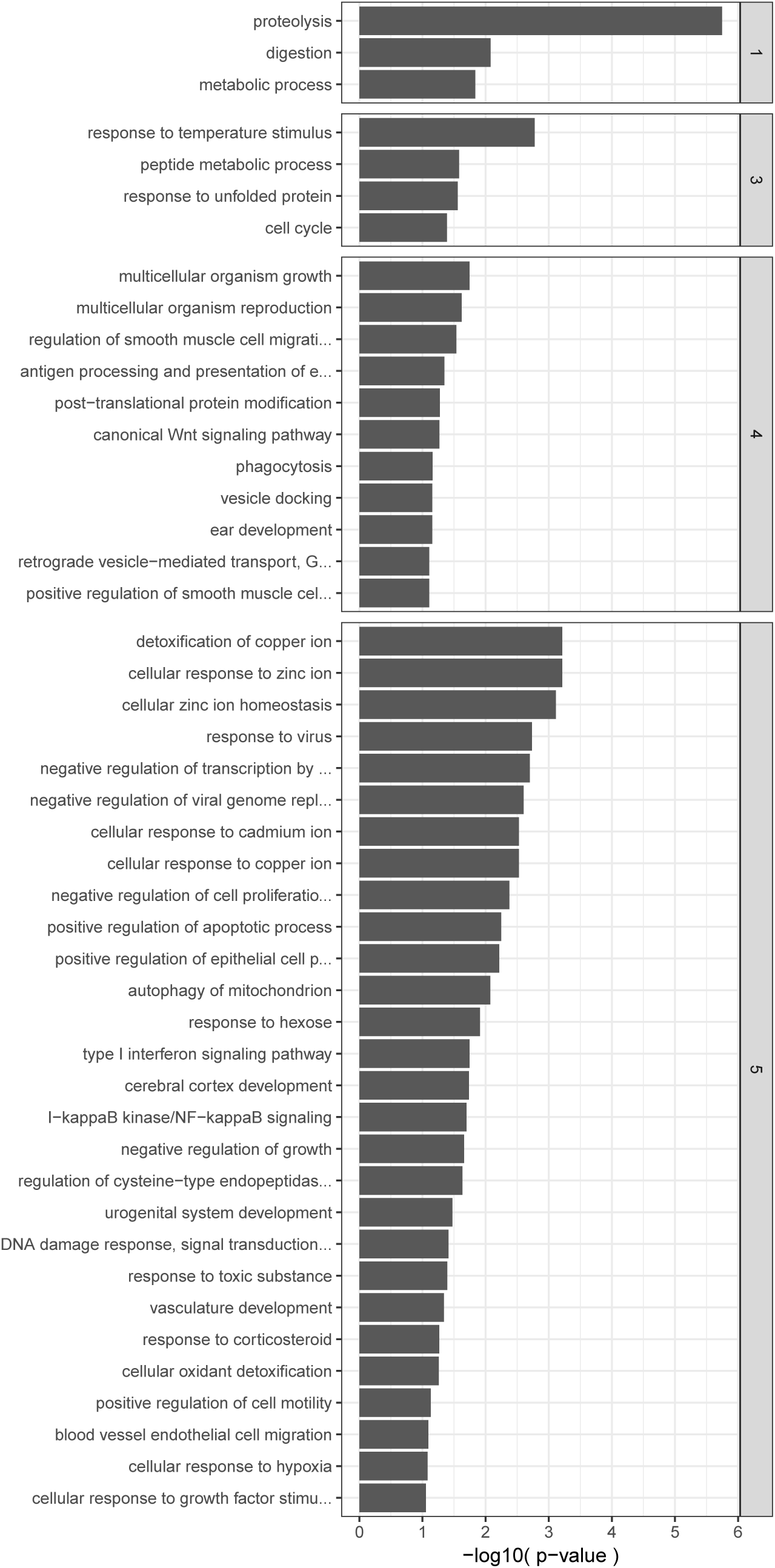
Enriched GO terms associated with gene clusters, ordered by psupertime. Biological process GO terms enriched in each gene cluster, relative to all other gene clusters. *x*-axis is uncorrected *p*-values for Fisher’s exact test. Only GO terms with at least 5 genes annotated in the cluster and *p*-value < 0.1 are shown (this results in no GO terms shown for cluster 2). See subsection 3.6 for details of calculation.

**Supp Fig 6:**
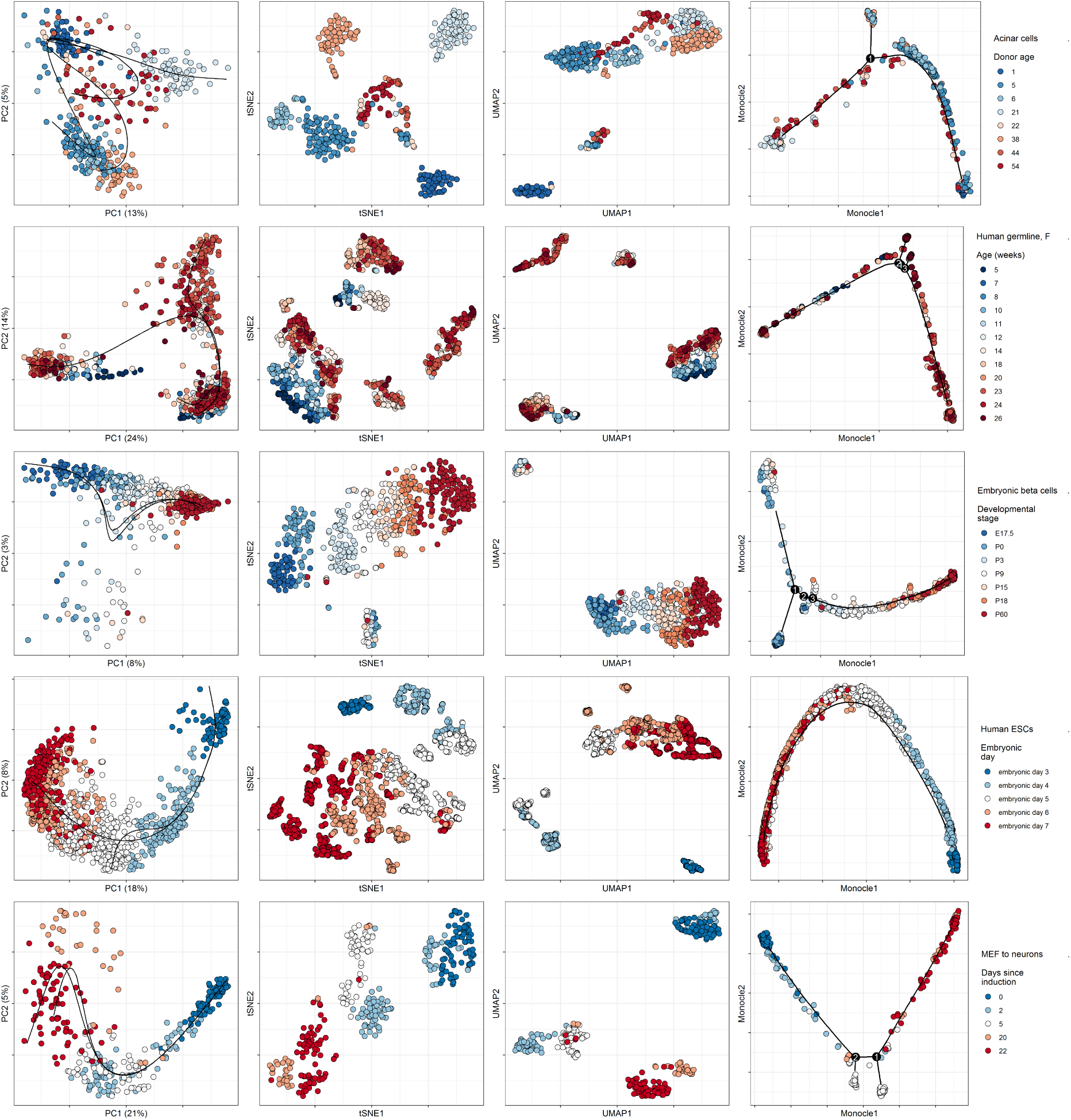
Dimensionality reduction methods applied to comparison datasets. Rows correspond to datasets detailed in Table 1. Columns correspond to dimensionality reduction methods, with annotations by comparator pseudotime inference techniques. First column corresponds to projection into the first two principal components, annotated with curves learned by `slingshot` [17], which are used as pseudotime. Second column shows projection by t-SNE, using default parameters [35]. Third column shows projection by `UMAP`, using default parameters [21]. Fourth column shows dimensionality reduction by `Monocle 2`, annotated with the tree it learns and which is used for pseudotime inference [16].

**Supp Fig 7:**
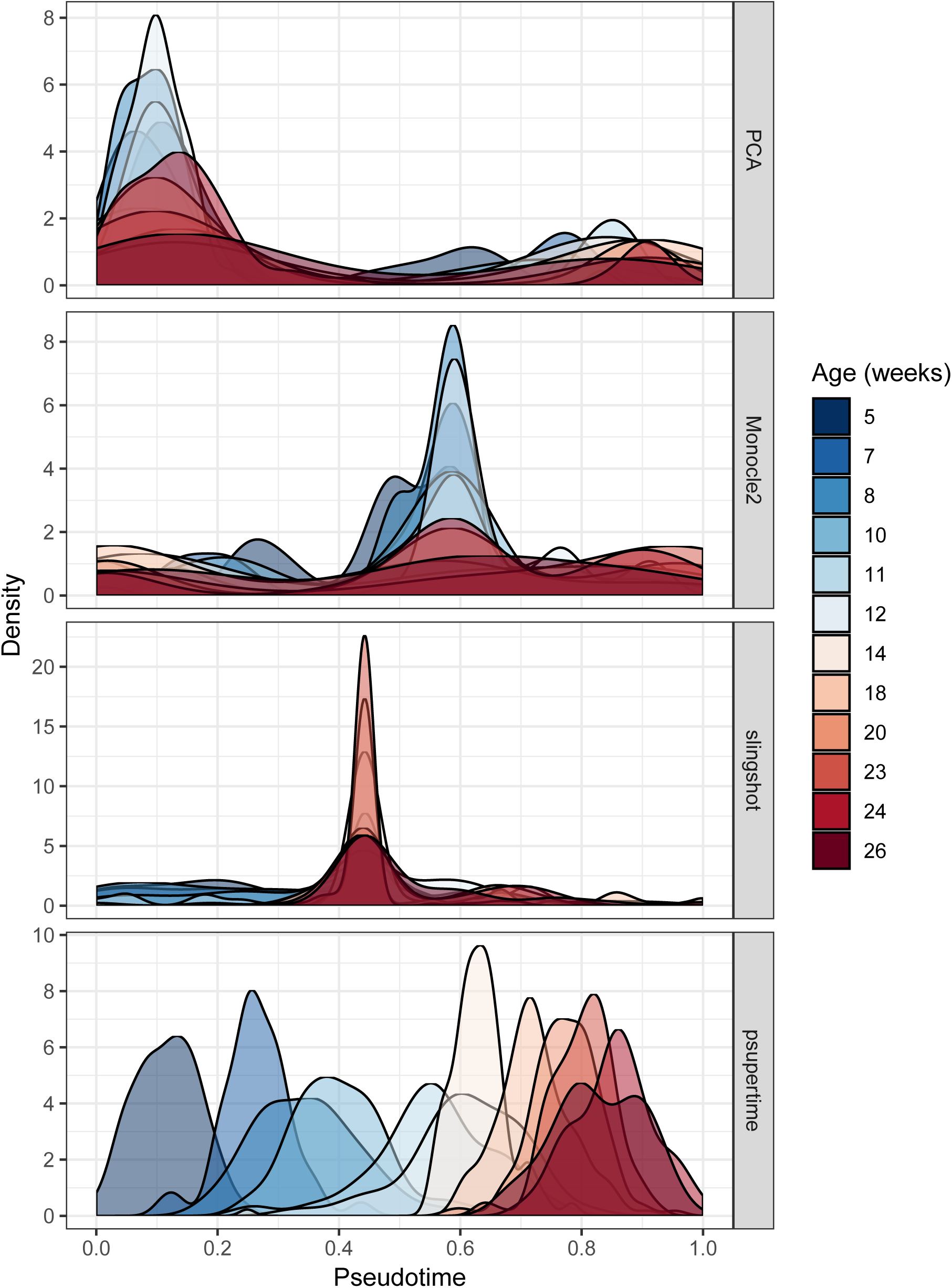
Benchmark methods applied to female human germline data. Female human germline data; colours indicate weeks post-fertilization. *x*-axes are the pseudotimes generated by each method, scaled to take values between 0 and 1. *y*-axes are density of the distributions for each label used as input, as calculated by the function `geom_density` in the `R` package `ggplot2` [23].

**Supp Fig 8:**
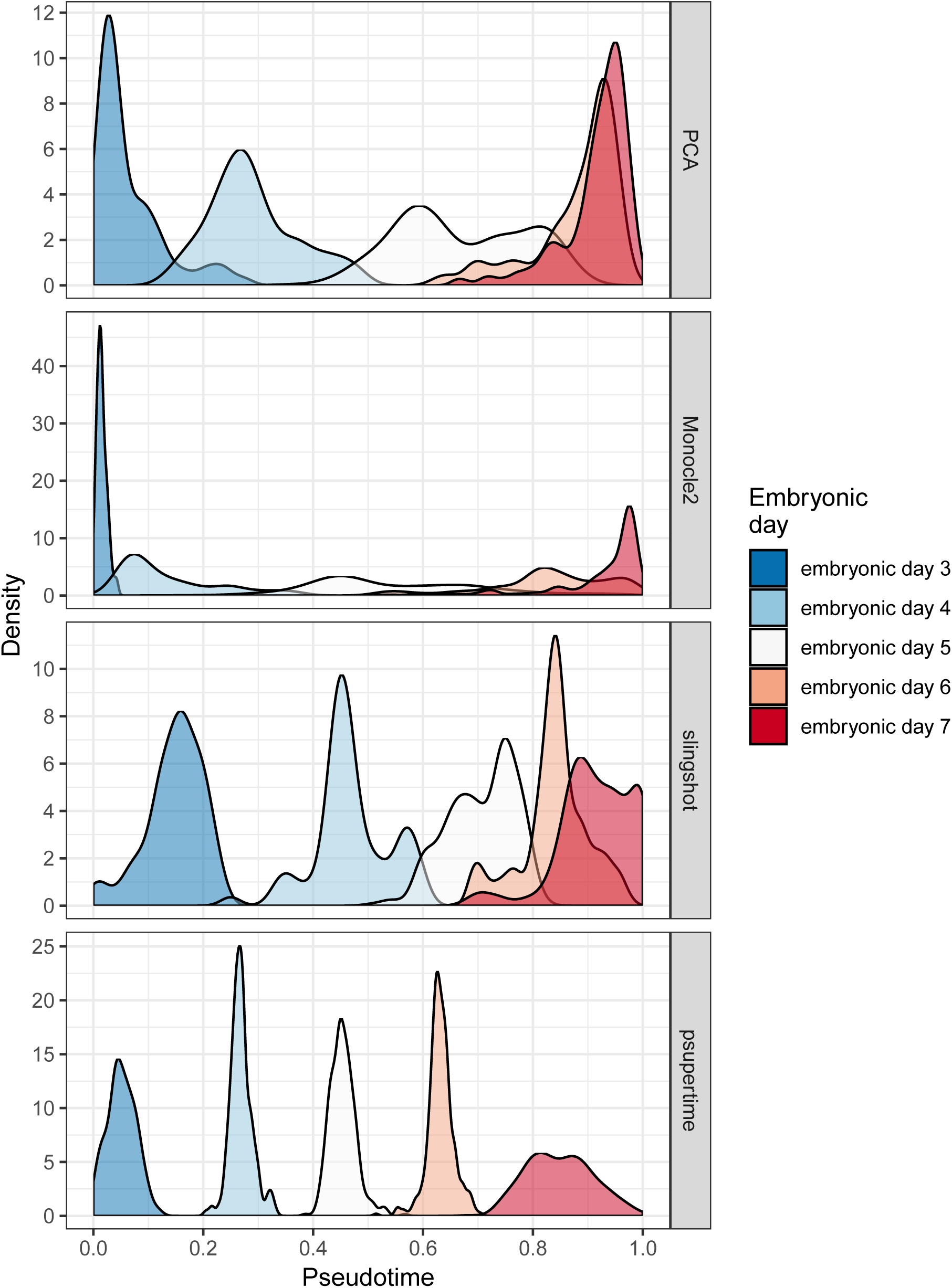
Benchmark methods applied to human ESC data. Human embryonic stem cell (ESC) data; colours indicate embryonic days. *x*-axis is the pseudotime generated by each method, scaled to take values between 0 and 1. *y*-axes are density of the distributions for each label used as input, as calculated by the function `geom_density` in the `R` package `ggplot2` [23].

**Supp Fig 9:**
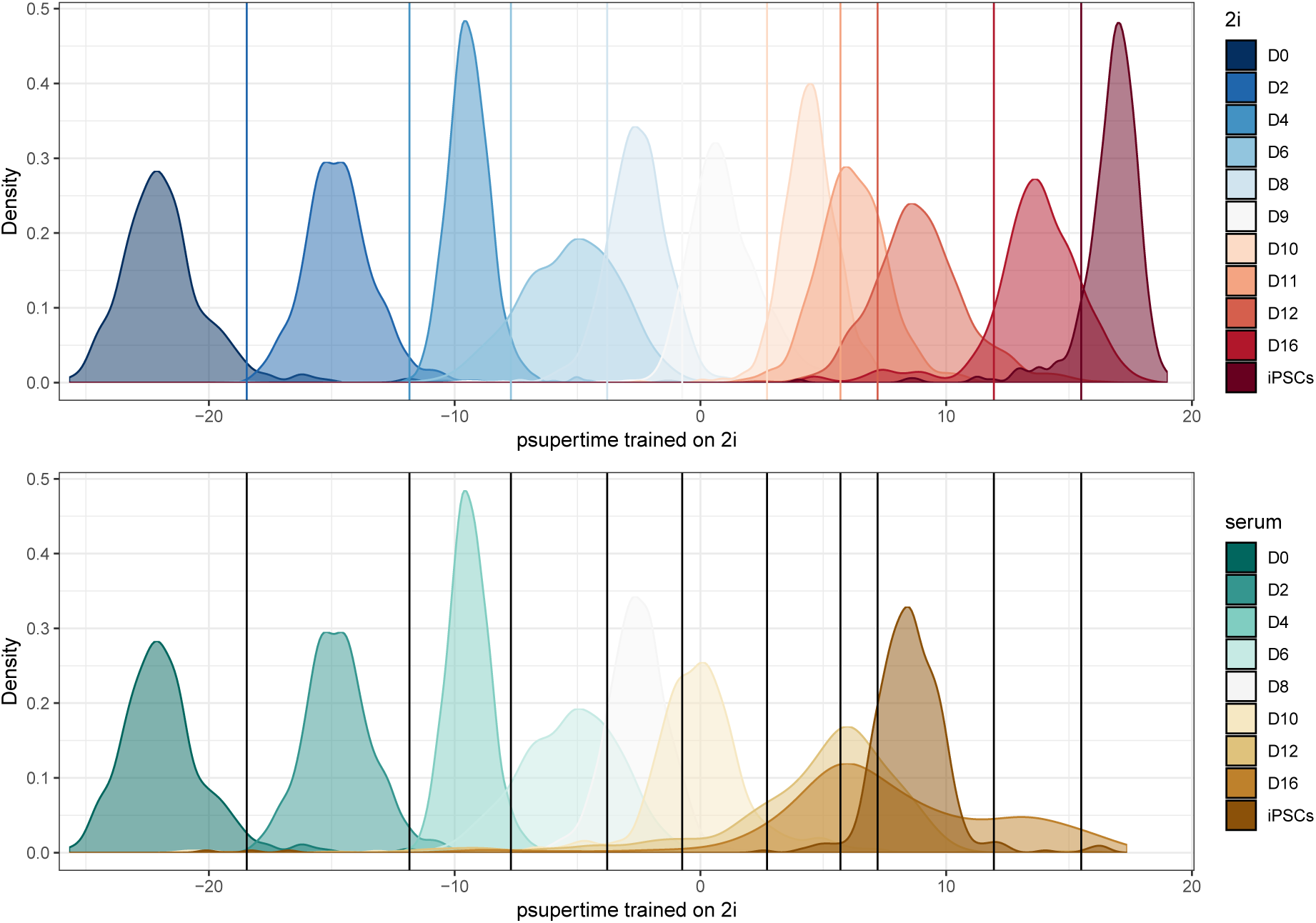
psupertime trained on 2i condition, used to predict labels for serum condition-pseudotime values. Data comprises: 3600 cells over labels D0 to D8 under DOX condition; 3600 over labels D9 to iPSCs under 2i condition; 1600 over labels D10 to iPSCs under serum condition [19]. `psupertime` was trained on cells taken from the DOX + 2i condition. The first row shows the distribution of the labels for these cells, and the corresponding pseudotime values from `psupertime`. The second row shows the results of applying this `psupertime` to cells from the DOX + serum condition. The labels are the true labels from this condition, and the 2i-trained `psupertime` was used to predict their *x*-axis values. For the cells over days D0 to D8, the cells used are identical, resulting in identical distributions over the pseudotime. On both plots, the vertical lines indicate thresholds between the labels for the 2i condition.

**Supp Fig 10:**
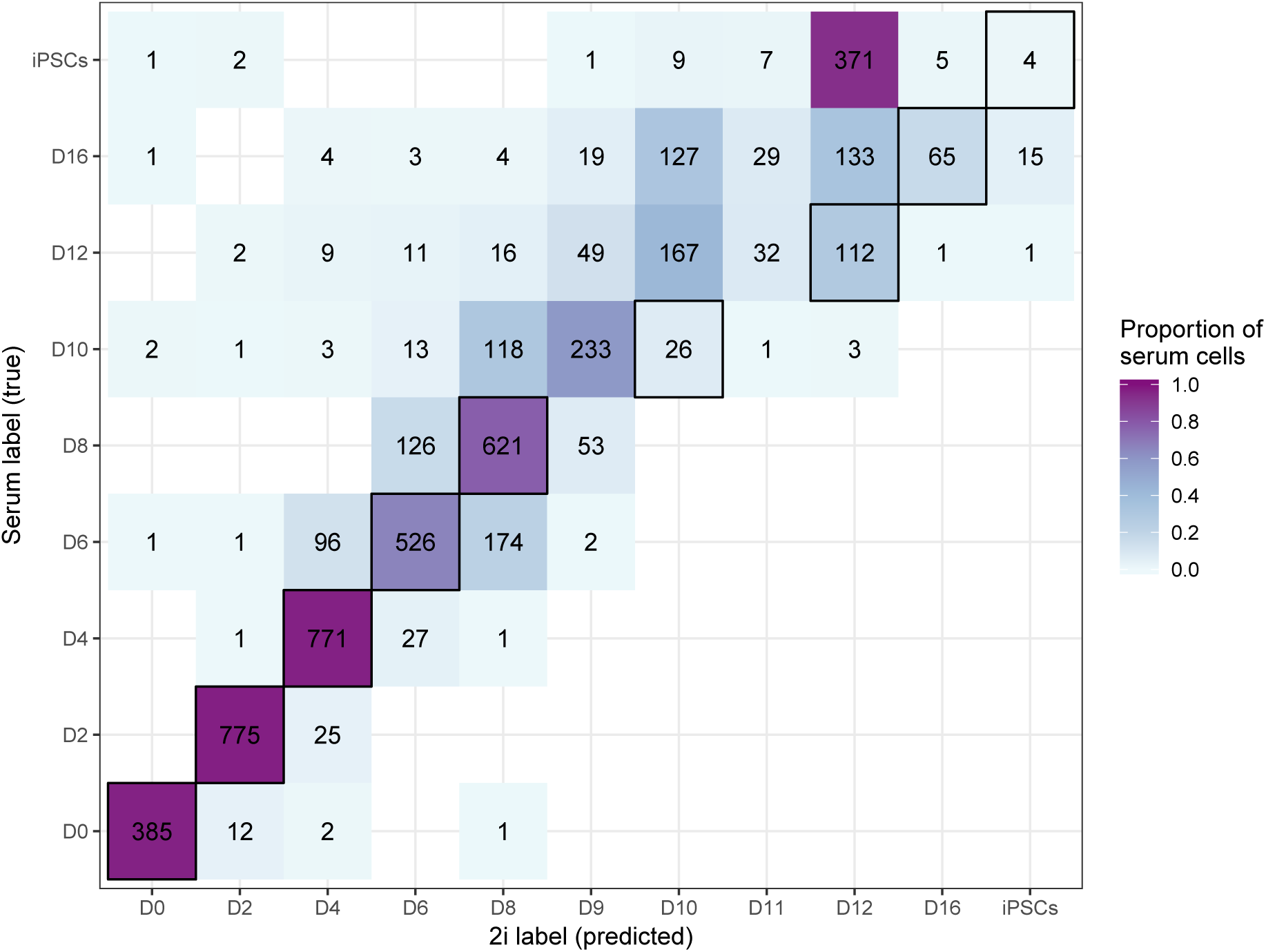
psupertime trained on 2i condition, used to predict labels for serum condition-confusion matrix. Data and application of `psupertime` as for Supp Fig 9. The rows show the true labels of cells from the DOX + serum condition. The columns show the predicted labels from the `psupertime` trained on cells taken from the DOX + 2i condition. Numbers correspond to number of cells with this combination of predicted and true labels.

**Supp Fig 11:**
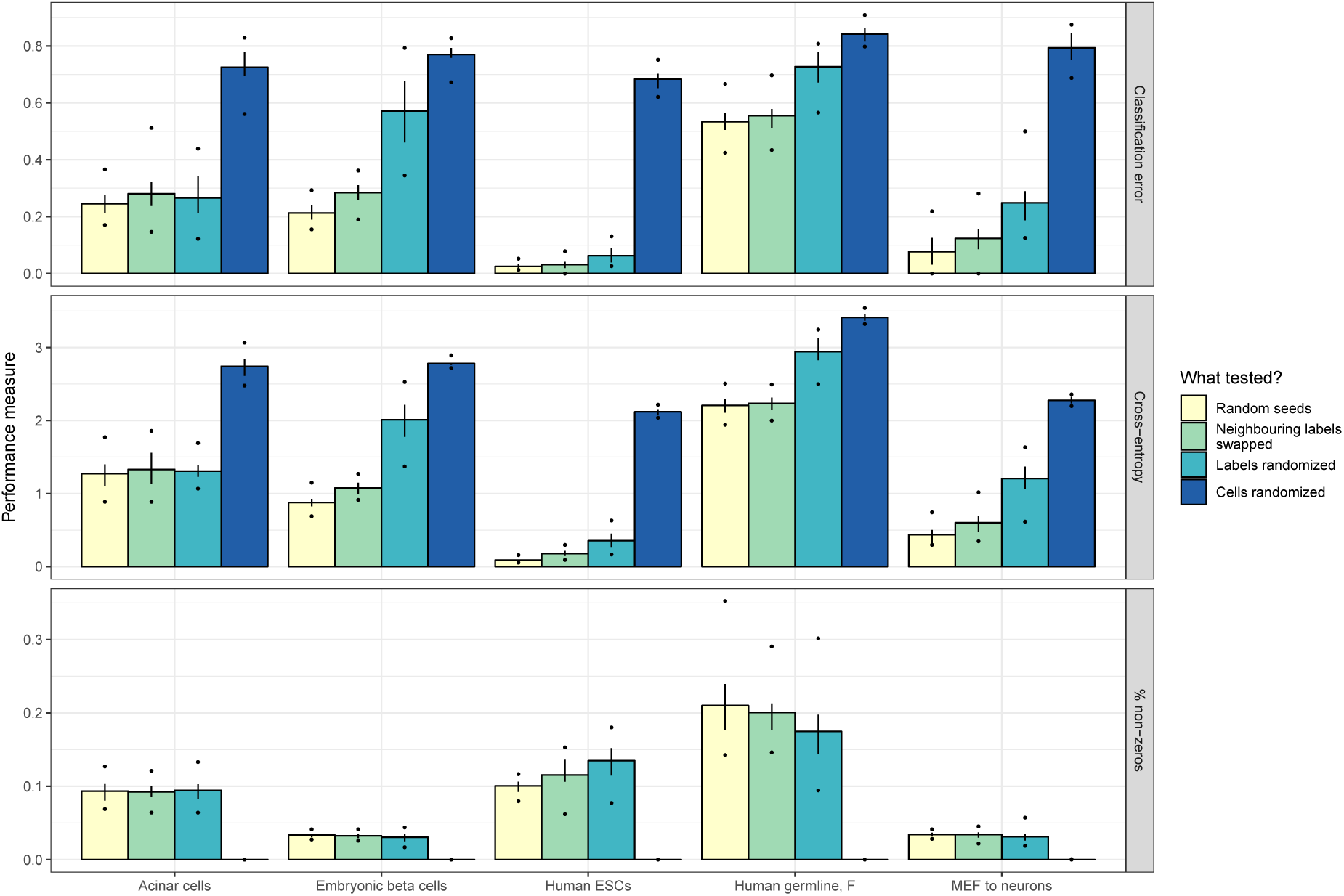
Robustness of psupertime to perturbations of labels. Rows correspond to measures of `psupertime`’s performance: classification error is the proportion of labels correctly assigned by `psupertime`; cross-entropy is a measure of confidence in predictions, which is high when a correct label is predicted with high probability; proportion of non-zero genes indicates what proportion of the input genes for which `psupertime` identified non-zero coefficients. Both classification error and cross-entropy are shown for test data, i.e. cells which were not used to train `psupertime`. The line range shows interquartile range, dots show minimum and maximum values observed, both over 20 random runs. The *x*-axis corresponds to the datasets detailed in Table 1.

**Supp Fig 12:**
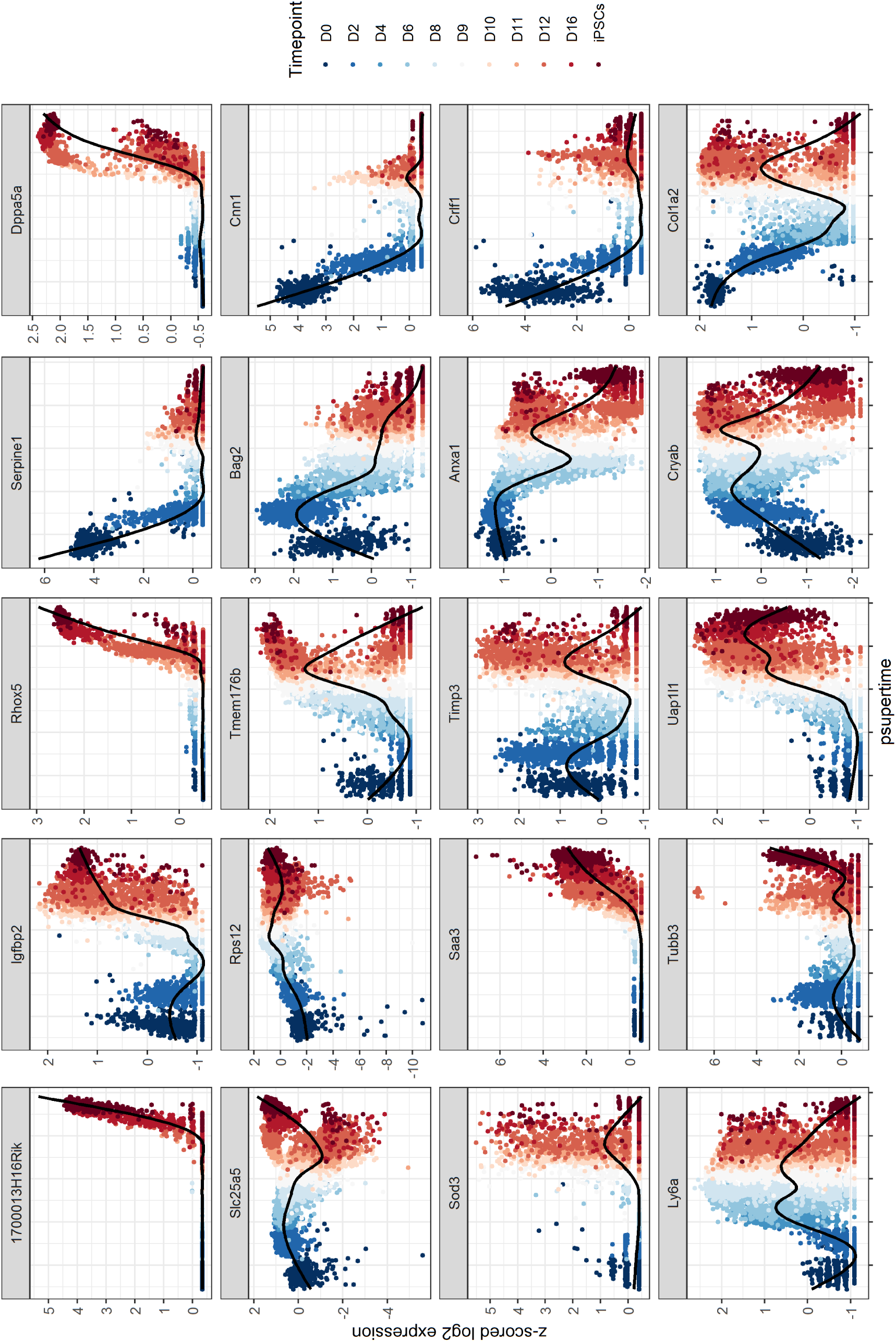
Profiles of top genes identified by psupertime in 2i-treated MEF cells. `psupertime` applied to 3600 cells over labels D0 to D8 under DOX condition; 3600 over labels D9 to iPSCs under 2i condition [19]. 20 genes with highest absolute coefficients, plotted against `psupertime` pseudotime. *x*-axis is the values from projections of each cell by `psupertime`. *y*-axis is smoothed, z-scored log pseudocounts for each cell. Colours indicate ordered labels. Black line is smoothed curve as fit by `geom_smooth` in the R package `ggplot2` [23].

**Supp Fig 13:**
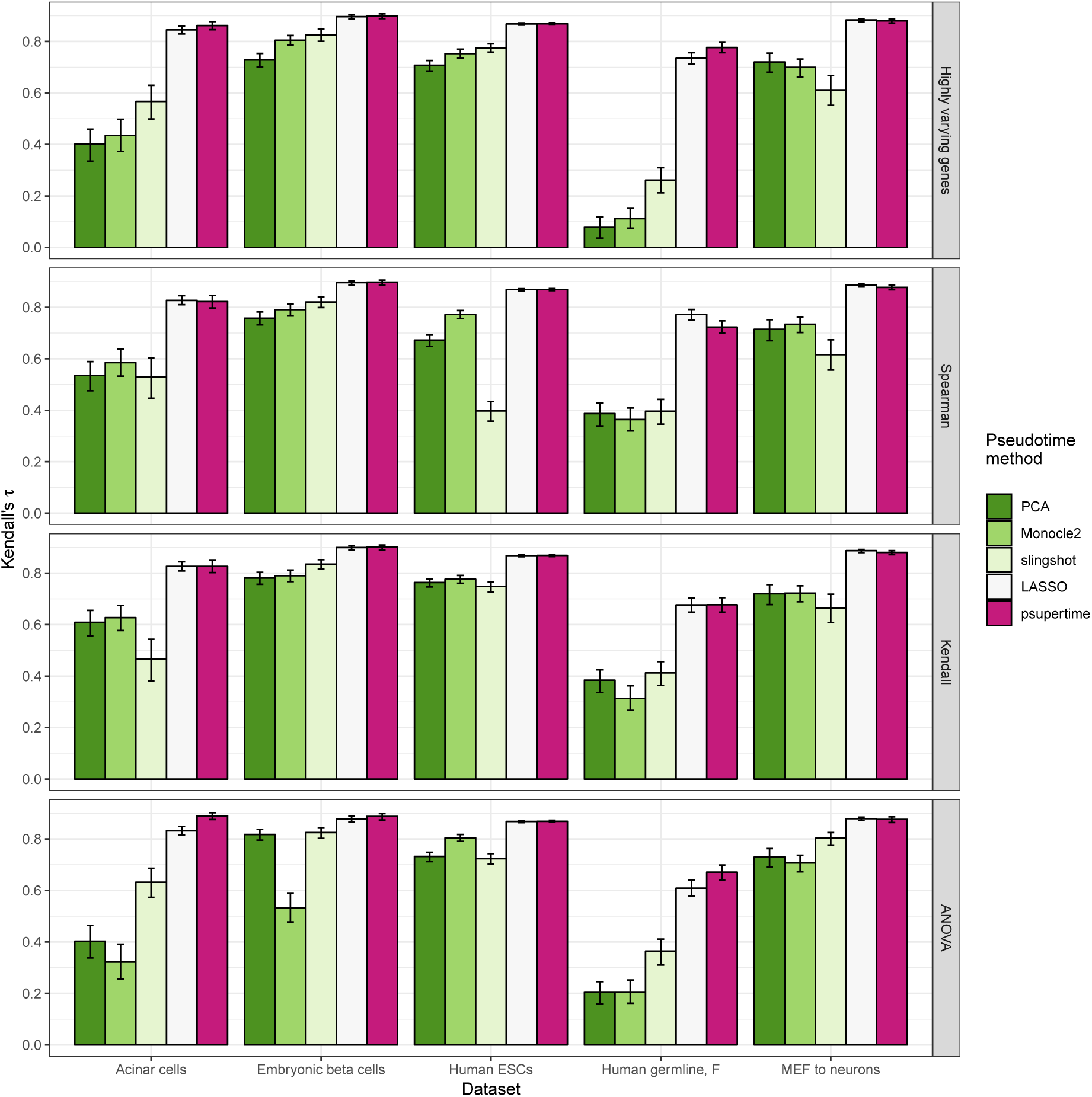
Performance of benchmark methods under different methods for gene selection. Rows correspond to different methods for selecting genes for input into the methods. The *x*-axis corresponds to datasets, detailed in Table 1. Colours indicate the benchmark pseudotime inference approaches described in subsection 3.5. The *y*-axis shows Kendall’s *τ* statistic, which assesses the extent of discordance between two orderings. For each combination of dataset and gene selection method, the five tested pseudotime approaches use exactly the same genes as inputs. Error bars show 95% confidence interval over 1000 bootstraps, calculated with `boot` package in R [24].

**Supp Fig 14:**
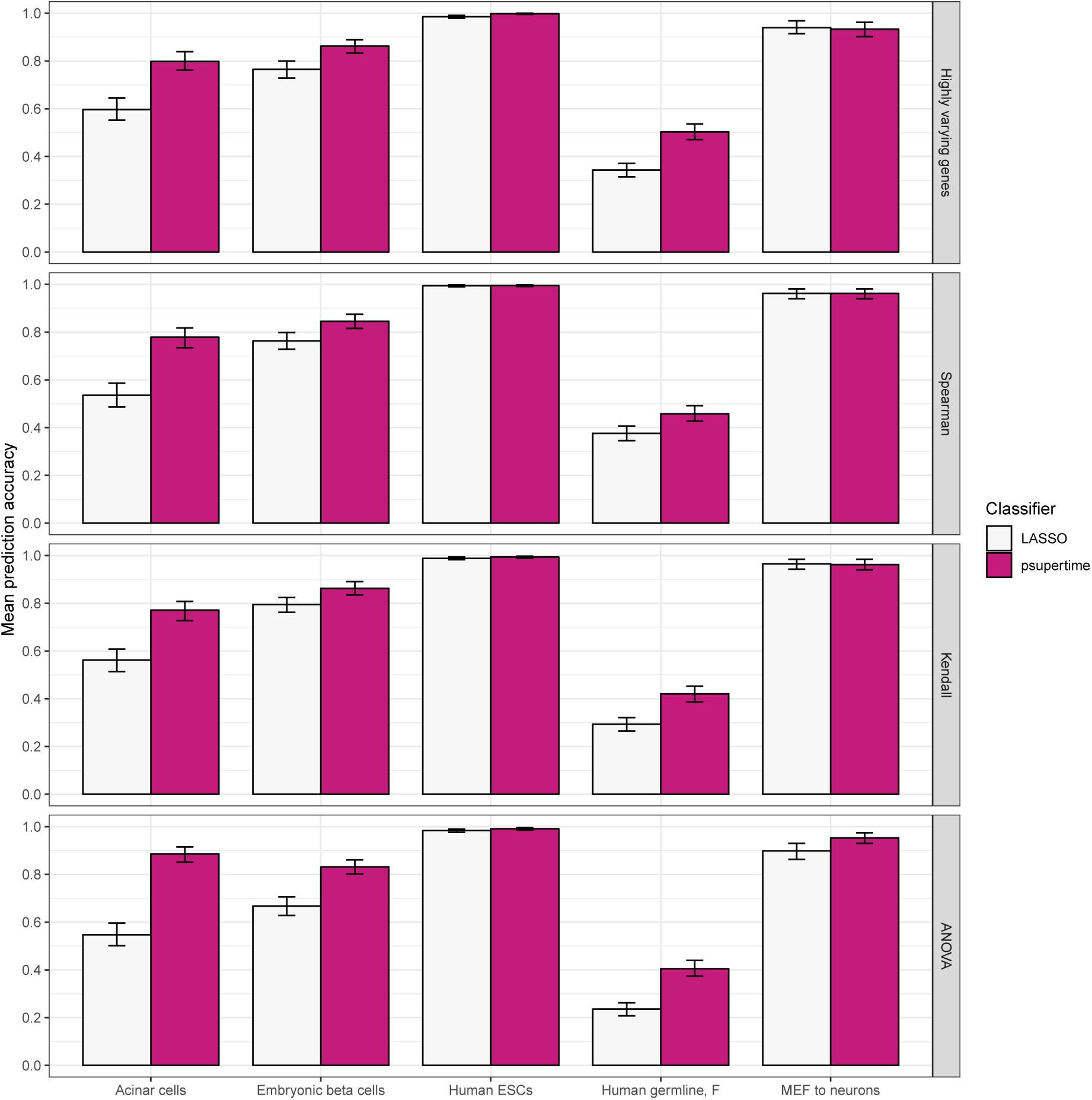
Classification performance of LASSO and psupertime under different methods for gene selection. Rows correspond to different methods for selecting genes for input into the methods. The *x*-axis corresponds to datasets, detailed in Table 1. *y*-axis shows the mean prediction accuracy over all cells. Classification under LASSO is done by fitting the model, calculating the estimated value ŷ for a given cell, and reporting the closest integer to ŷ in 1, …, *K*. For each combination of dataset and gene selection method, the two tested classifiers use exactly the same genes as inputs. Error bars show 95% confidence interval over 1000 bootstraps, calculated with boot package in R [24].

**Supp Fig 15:**
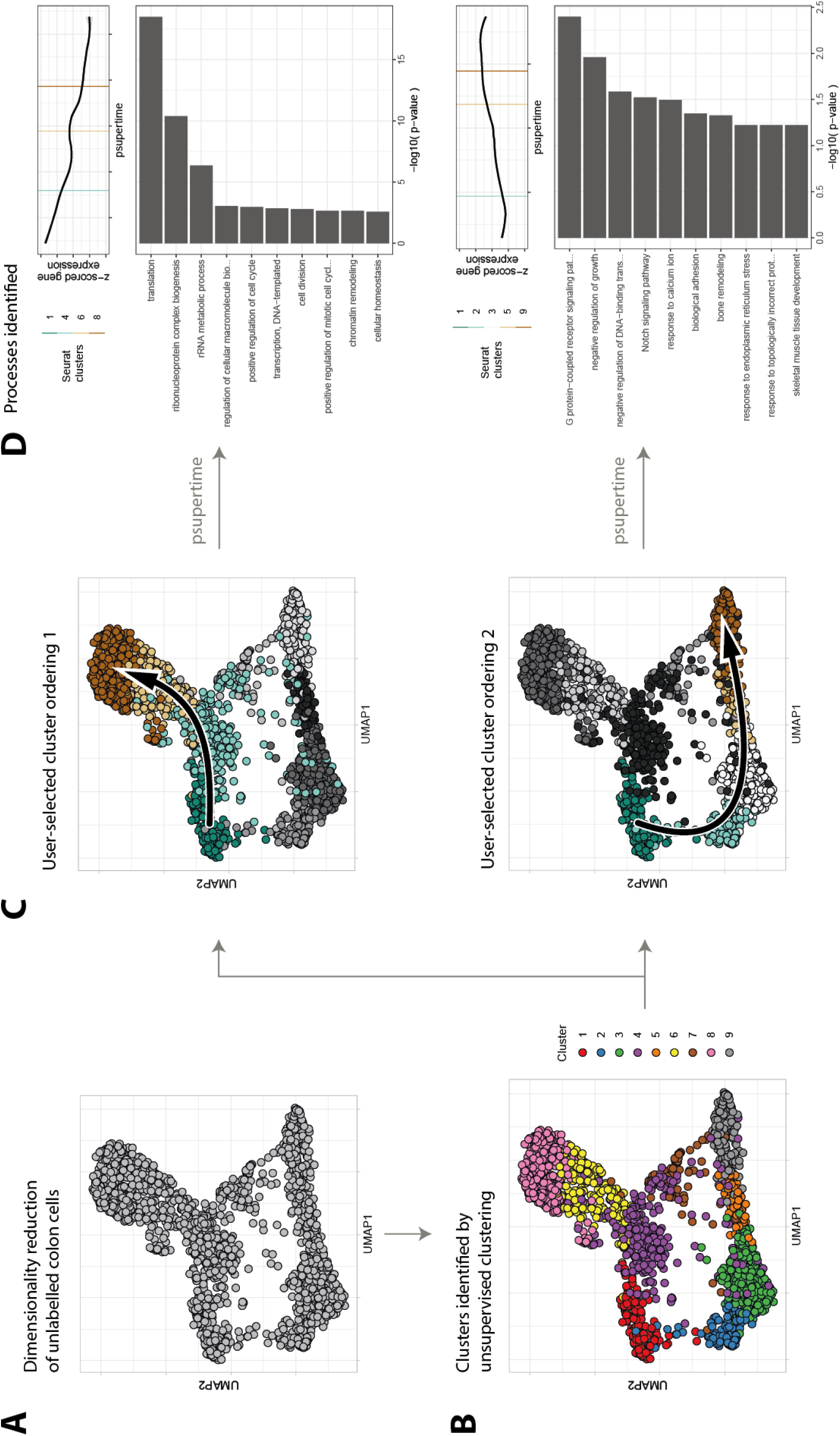
psupertime used for exploratory data analysis. A Dimensionality reduction (UMAP [21]) of 1894 colon cells [14]. **B** Unsupervised clustering (via R package Seurat [15]) identifies 9 clusters within the sample. **C** Users can select cluster sequences they wish to investigate; two are shown here, which may correspond to development from stem cells into distinct mature celltypes. Arrows indicate the selected sequence. **D** Geneset enrichment of clustered gene profiles identifies biological processes associated with the sequence. Hierarchical clustering identified 5 gene clusters; clusters shown here are those with highest positive correlation with learned pseudotime. GO terms shown correspond to the smallest 10 *p*-values, subject to *p <* 10% and at least 5 annotated genes.

**Supp Fig 16:**
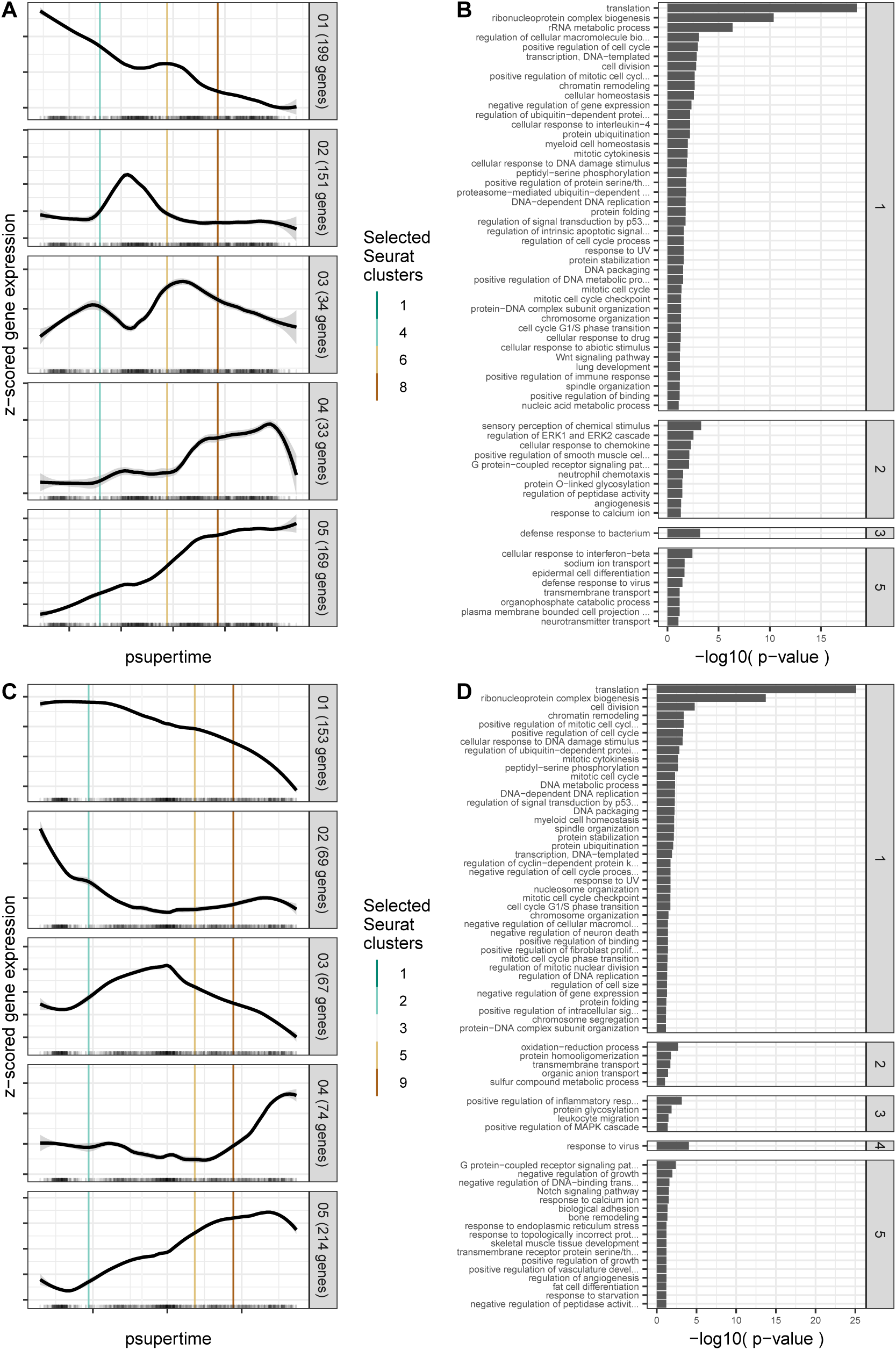
Biological processes associated with gene profiles identified by psupertime. Data comprises 1894 cells from colon [14]. `psupertime` was applied to two different user-selected cluster sequences. For each sequence, all genes selected for training were clustered into five clusters, and geneset enrichment analysis was used to identify biological processes distinctive of each cluster relative to the other four. See subsection 3.6 for details. **A** Clusters identified for cluster sequence 1468, ordered by correlation between mean profile and pseudotime. **B** Biological process GO terms identified as enriched in each cluster, relative to the remaining clusters; all GO terms shown have both Fisher exact *p*-value < 10% and at least 5 genes in the cluster annotated. **C, D** As for A,B but for cluster sequence 12359.

**Supp Fig 17:**
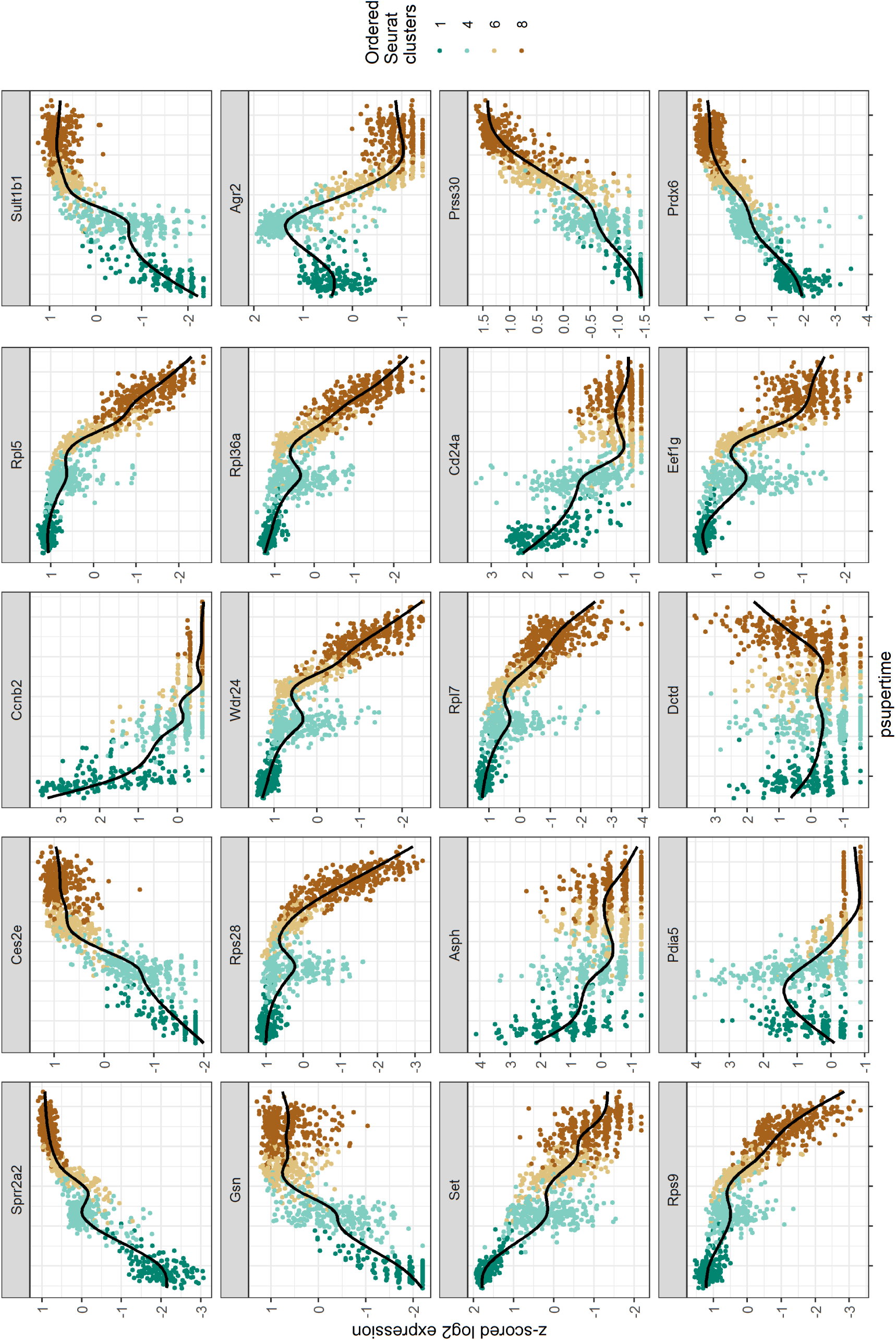
Profiles of top genes identified by psupertime for user-selected cluster sequence 1. 20 genes with highest absolute coefficients, plotted against `psupertime` pseudotime. *x*-axis is the values from projections of each cell by `psupertime`. *y*-axis is smoothed, z-scored log pseudocounts for each cell. Colours indicate ordered labels. Black line is smoothed curve as fit by `geom_smooth` in the R package `ggplot2` [23].

**Supp Fig 18:**
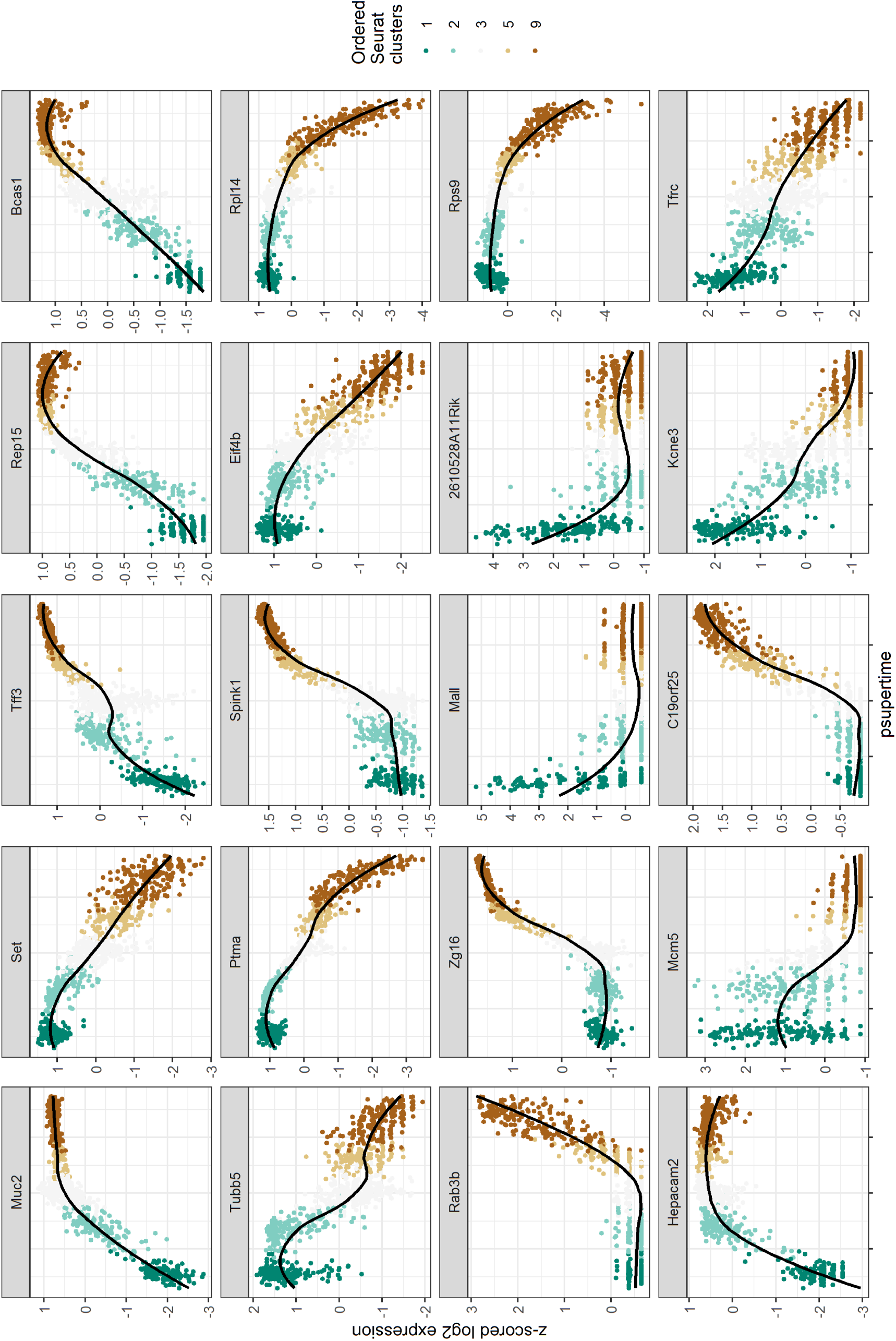
Profiles of top genes identified by psupertime for user-selected cluster sequence 2. 20 genes with highest absolute coefficients, plotted against `psupertime` pseudotime. *x*-axis is the values from projections of each cell by `psupertime`. *y*-axis is smoothed, z-scored log pseudocounts for each cell. Colours indicate ordered labels. Black line is smoothed curve as fit by `geom_smooth` in the R package `ggplot2` [23].

**Supp Fig 19:**
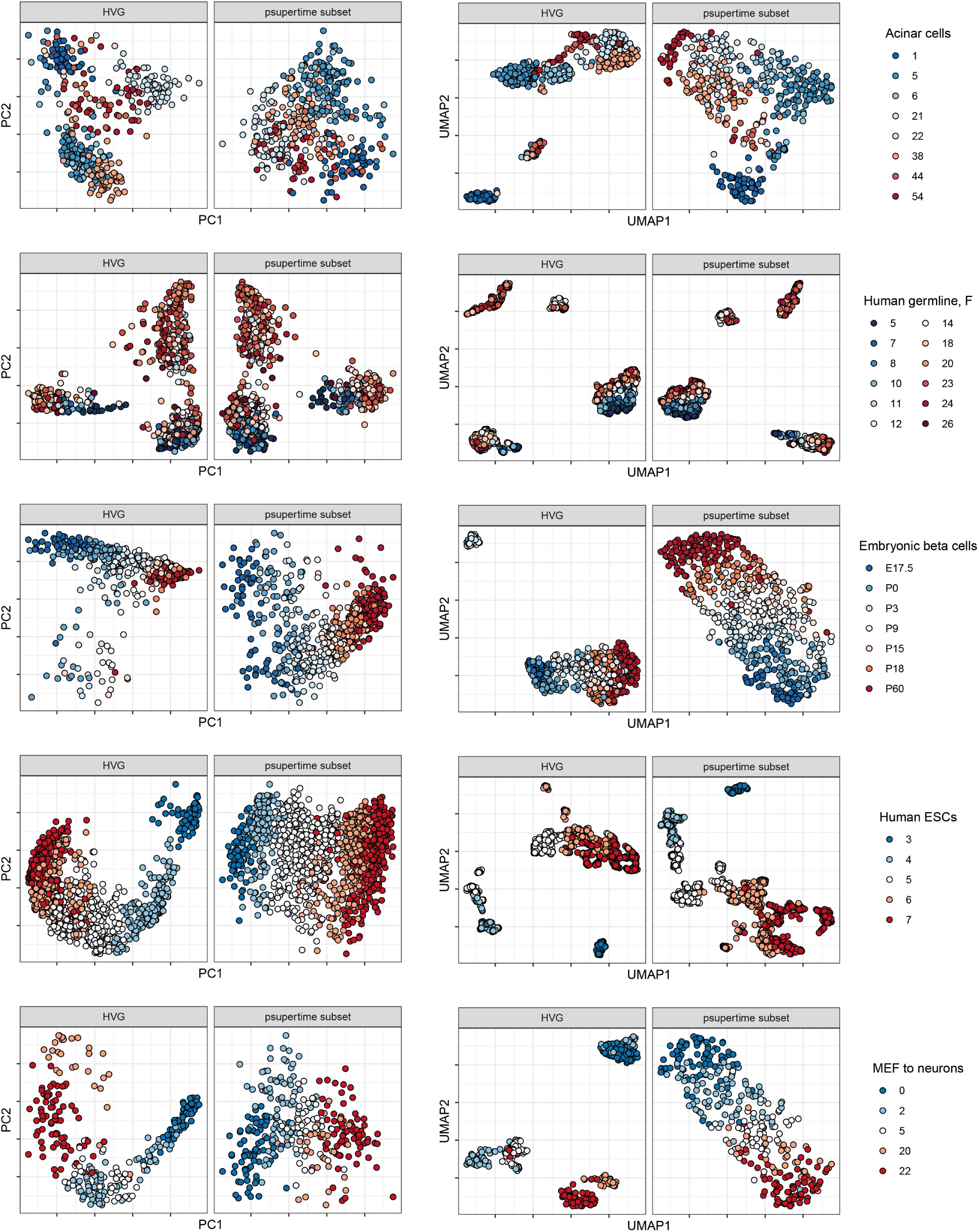
Dimensionality reduction comparing highly variable genes and genes identified by psupertime as input. Rows correspond to datasets detailed in Table 1. First pair of columns shows first two principal components; second pair shows projection with UMAP, using default parameters [21]. In both pairs of columns, the left uses all highly variable genes (‘HVG’) as input, and the right uses only genes identified by `psupertime` as input (‘`psupertime` subset’).

